# The Phenotypic Expression of Purple Body (*Pb*) in Domestic Guppy Strains of *Poecilia reticulata*

**DOI:** 10.1101/121301

**Authors:** Alan S. Bias, Richard D. Squire

## Abstract

Modification of wild-type carotenoid orange and pteridine red coloration and spotting of male ornaments in modern Domestic Guppy Strains (*Poecilia reticulata reticulata*) by the naturally occurring Purple Body gene (Pb) has been long incorporated into their strains by Pedigree Stock Breeders. It is inherited as an autosomal incompletely dominant trait. Its existence has allowed breeders to produce a vast array of Purple based phenotypes. Photographic evidence demonstrates that Purple Body is a normal polymorphism in domestic guppies modifying color pigmented regions. When combined with currently used mutant genes such as Albino, Blond, Golden, Asian Blau, Coral Red, Magenta, Grass, Moscow, Pink, Platinum, Red Mosaic, Multicolor, and Full Red, startling new phenotypes are created. The recently described Purple Body gene (Bias and Squire 2017a, 2017b, and 2017c) has long been overlooked in research articles and little understood in breeder publications.

## Introduction

The Purple Body gene is located on an autosome. Breeding tests, involving this modification of orange spotting, reveal this trait to have an incompletely dominant mode of inheritance. As such a formal name and nomenclature of **Purple Body (*Pb*)** has been suggested (Bias and Squire, 2017a). *[**Note**: Hereafter Purple Gene, or Purple Body Gene used interchangeably in reference.]*

The intent of this brief paper is to provide spectral distinction, for ease in identification, between Pb and non-Pb in many of the more commonly produced modern Domestic Guppy strains. Emphasis is on the primary color traits, and several common pattern traits. Groupings are presented by Grey (*wild-type*), autosomal and sex-linked modifiers. These should not be considered inclusive of all known phenotypes. The number of strains and phenotypes being produced for color and pattern continues to increase, and is a testament to the efforts of both professional and amateur breeders around the world.

Autosomal incompletely dominant Pb, similar to identified autosomal recessives found in *Poecilia reticulata*, is a modifier of total existing body color and pattern pigmentation (involving xantho-erythrophores, structural iridophores and melanophores) in both males and females. Purple Body is capable of modifying extent color and pattern found in any Domestic Guppy strain.

Pb modification is most noticeable in Domestic Strains as a modifier of ornaments comprised of orange carotenoid color pigment, in both males and females **(Fig 2)**. Visually, coloration is modified from a highly reflective orange to a “pinkish-purple” coloration in Grey (“wild type” alleles *A, B, G, R*, and *ab*) corresponding to autosomal genes Albino (*a*, Haskins and Haskins 1948), Blond (*b*, Goodrich 1944) Golden (*g*, Goodrich 1944), European Blau (*r*, Dzwillo 1959) Asian Blau (*Ab, Undescribed* - see Bias 2015). It is also modified in various ways when combined with autosomal genes Pink (*p*, Luckman 1990, Förster 1993, *pi*, Kempkes 2007), Ivory (*I*, Tsutsui 1997, Magenta (*M*, undescribed), and Zebrinus (*Ze*, Winge 1927). It combines as well with sex-linked genes such as Coral Red (*Co, undescribed*) Y-linked, Grass (*Gra, undescribed*) X- and/or Y-linked, Moscow (*Mw*, Y-linked, Kempkes 2007]), Nigrocaudautus, X and/or Y-linked (*NiI*, Nybelin 1947] and *NiII*, Dzwillo 1959). Platinum (*P, undescribed*) X- and/or Y-linked, Mosaic (*Mo*, Khoo and Phang 1999) X- and/or Y-linked, Multicolor (no gene symbol) X-linked, and Schimmelpennig Platinum (*Sc*, described as Buxeus by Kempkes 2007), Y-linked. Examples provided in this paper are primarily limited to Delta Tail and Swordtail phenotypes.

In heterozygous condition (*Pb/pb*) a distinct result is generated while in homozygous condition (*Pb/Pb*) these results are further amplified. Pb is capable of pleiotropic effect on all existing color and pattern elements at multiple loci. The purple phenotype has been present in hobbyist stocks for decades, but has been largely unrecognized by many breeders, except in the case of pure-bred all-purple strains.

Pb modification, zygosity dependent, removes certain classes of yellow-orange-red color pigment over silver iridophores or white leucophores. Pb modifies “other existing” color in both body and fins, thus suggestive of being a “full body” modifier, in homozygous condition. Dark red pteridine color pigment does not seem to be modified by Purple Body in fins lacking an underlying silver iridophore or white leucophore pattern. Modification by Pb seems limited predominantly to wild-type orange color pigment; i.e. that which also contains yellow carotenoids in addition to red pteridines, over an iridophore pattern.

Pb is always found in all-purple fish, but is not by itself sufficient to produce the all-purple phenotype in heterozygous expression. Homozygous Pb expression results in the further removal of xantho-erythrophores, in conjunction with both increased populations and/or greater visibility of modified melanophores and naturally occurring violet and blue iridophores. It is required for the production of the all-purple phenotype **(Fig 1 and Fig 2A)**. Pb causes a large reduction of yellow color pigment cell populations (xanthophores). It thus produces a modified pinkish-purple expression from what would have been orange color pigment cells (xantho-erythrophores).

**Fig 1.**
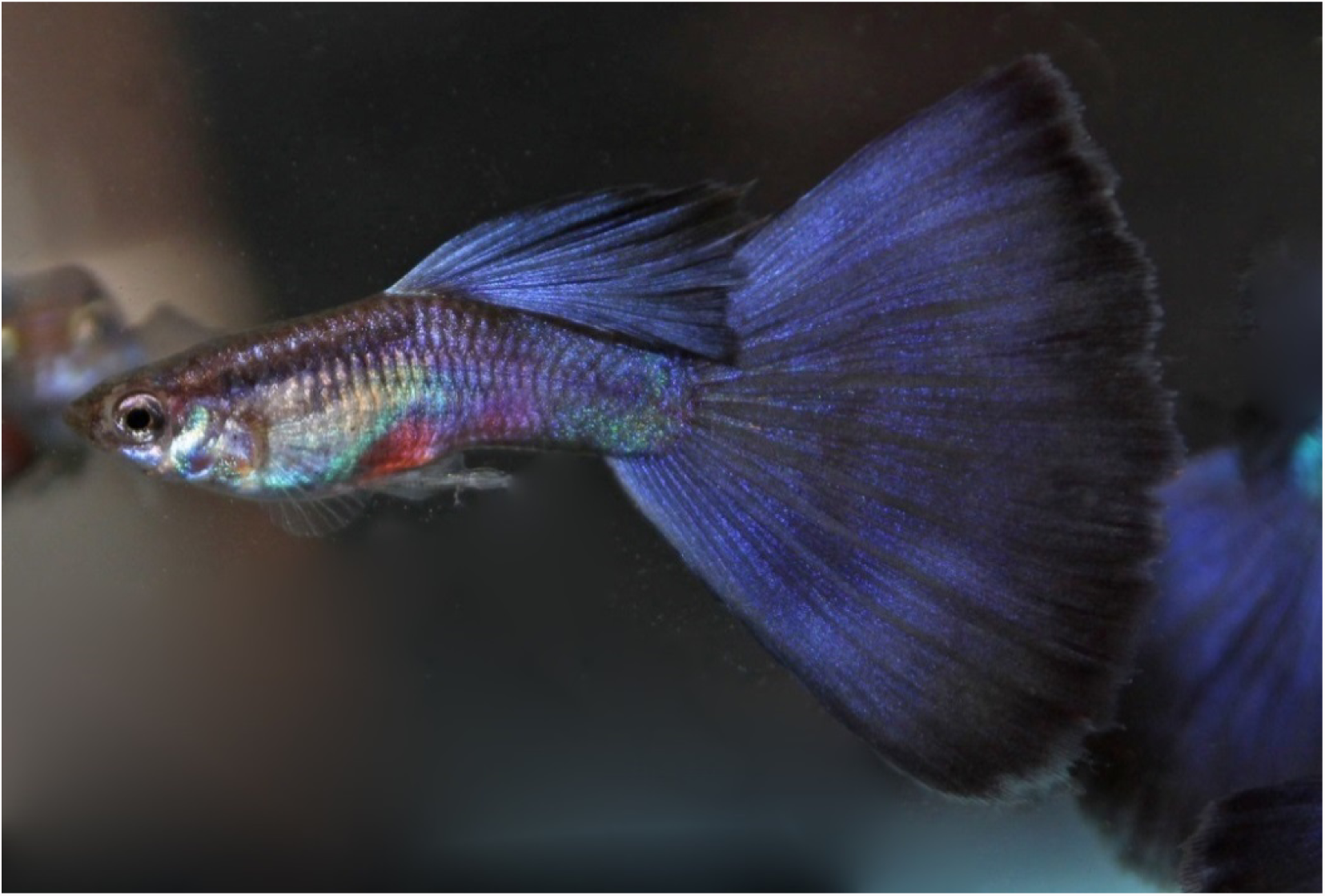
Purple Delta (*Pb/Pb*), photo courtesy of Terry Alley

**Fig 2.**
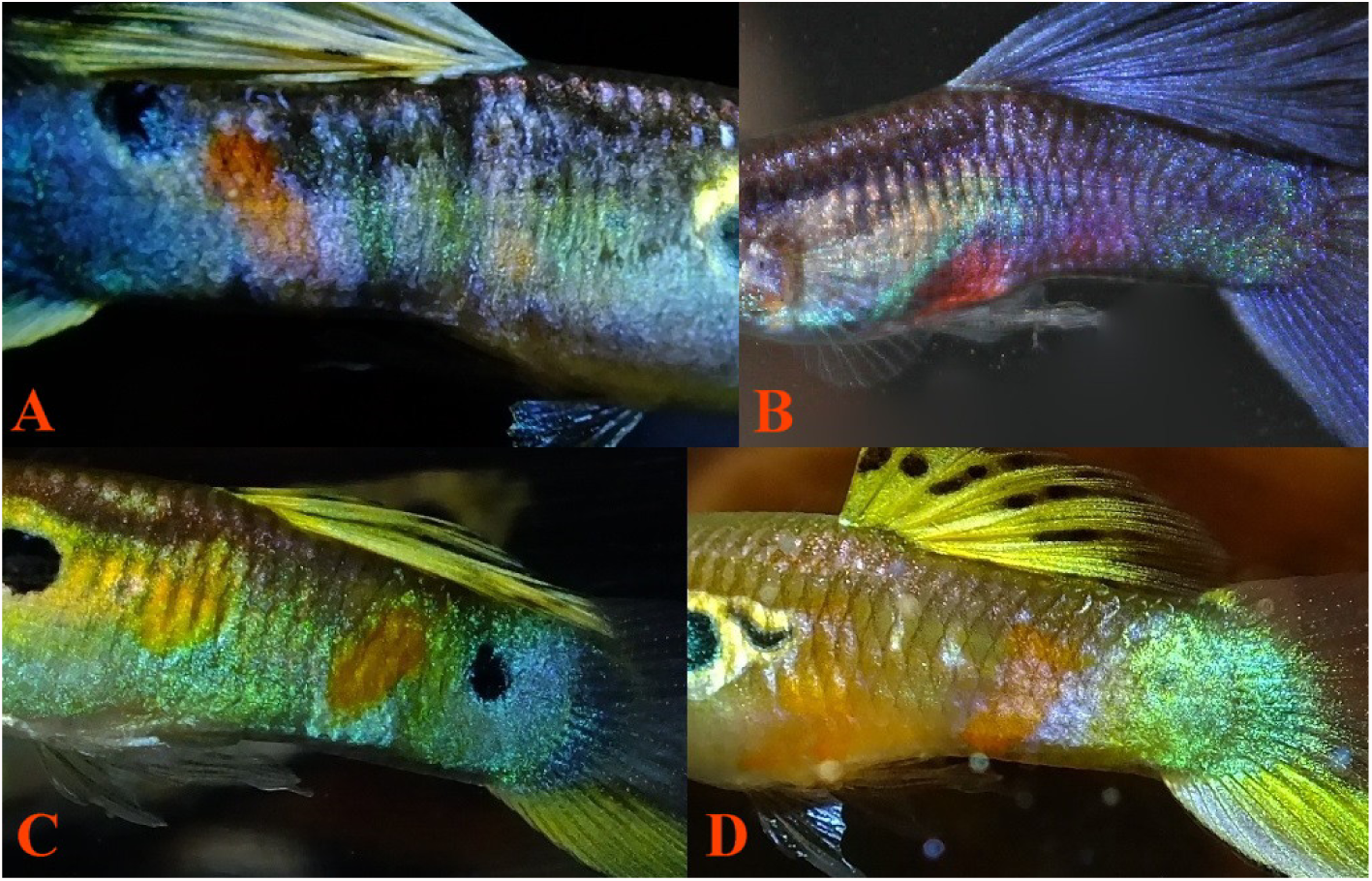
**(A)** Homozygous Pb (*Pb/Pb*) modified ornaments, expressing removal of xanthophores and increased violet-blue iridophores. **(B)** Homozygous Pb (*Pb/Pb*) modified ornaments, expressing reduced xanthophores and increased violet-blue iridophores. **(C-D)** non-Pb ornaments (*pb/pb*) expressing no alteration of xantho-erythrophores.

High resolution photography and microscopic study shows the co-existence of varying populations of both violet and blue structural iridophores in all individuals, both male and female (Bias and Squire 2017a, Pb Cellular Description; 2017b, Pb Microscopy Study; 2017c, Ocular Study). Violet and blue structural iridophores and melanophores are always found in close proximity with one another, forming a type of chromatophore unit [***Note**: hereafter referenced as violet-blue (iridophores) for ease of discussion*]. Violet-blue iridophores **(Fig 3A-B)** are most visible along the topline and in between regions lacking a clearly defined silver iridophore pattern, often including the caudal-peduncle base. By nature, yellow color pigment in Guppies is highly motile and mood dependent while red color pigment is considered non-motile. Red color pigment (from erythrophores) is not altered by Pb, or at least altered to a lesser degree, and a corresponding noticeable increase in the visibility (possibly increased population levels) of structural violet and blue iridophores is evident **(Fig 2A-B and Fig 3A-B)**, resulting in the increased reflective qualities of individuals.

**Fig 3.**
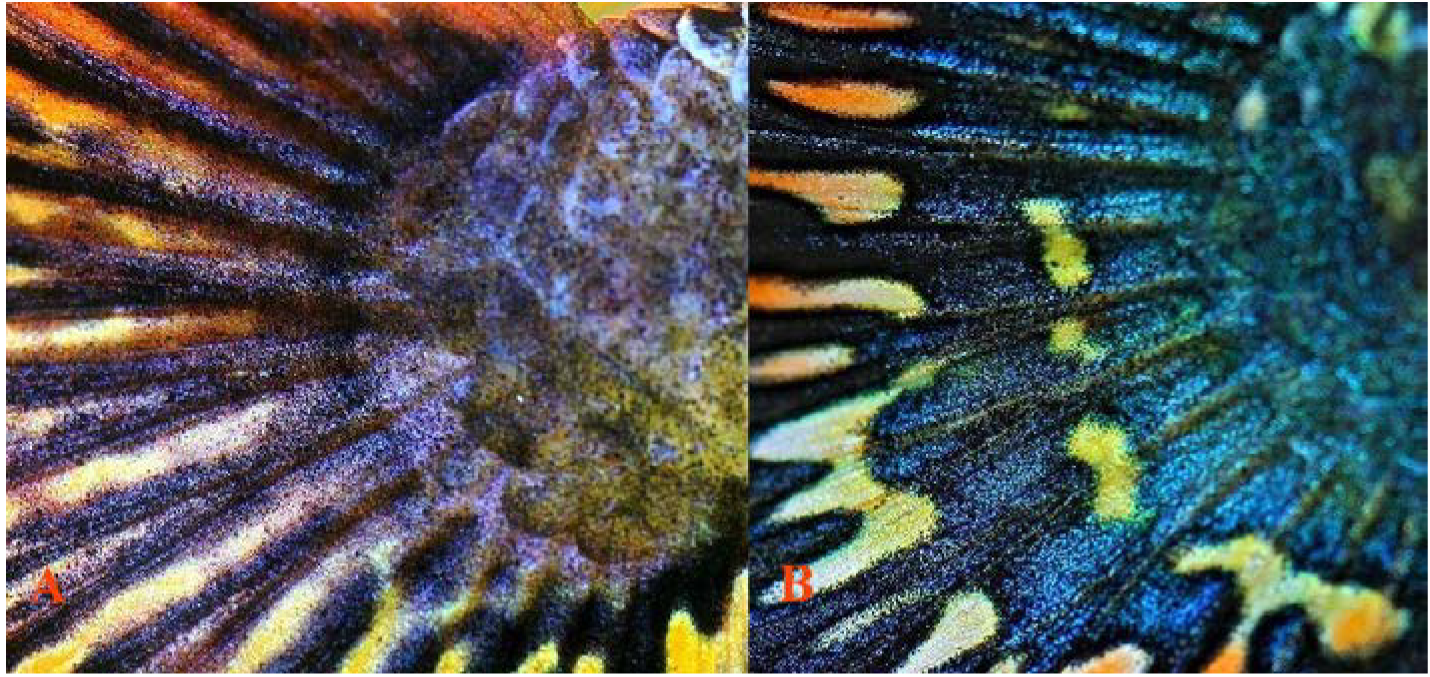
**(A)** Pb/Pb modified caudal base expressing increased violet iridophores. **(B)** Non-Pb pb/pb caudal base expressing balanced violet-blue iridophores, photos courtesy of Christian Lukhaup.

When not masked by additional color and/or pattern traits, the identification of Purple Body (*Pb*) in both wild-type and domestic males can be easily accomplished through visual phenotypic observation. In non-Purple Body (*pb/pb*) individuals carotenoid orange color pigment can be described as being vivid, bright orange spots structurally comprised of densely packed yellow and orange xantho-erythrophores, normally extending to the very edge of the spot. Though coverage over additional iridophore patterns may appear incomplete.

Heterozygous Purple Body (*Pb/pb*) alters orange spots in select regions of the body and in finnage to “pinkish-purple”. Thus, it may not act as a “full body” modifier in heterozygous form. Heterozygous Pb does not appear to greatly reduce visible structural yellow color pigment cells over white leucophore or reflective clustered yellow cells, known in breeder circles as Metal Gold (Mg) (*Undescribed* - Bias 2015), in body and finnage. A slight increase in visibility of violet and blue iridophores is often detected. Additionally noted is an increase and modification in existing melanophore structure and possibly population numbers, as compared to heterozygous Pb. In non-solid colored strains, a reduction in the number of yellow xanthophores results in a corresponding reduction in overall size of individual spotting ornaments. This reveals a “circular ring” around remaining color pigment produced by an underlying iridophore layer. This well-defined layer of iridophores is an underlying precursor required for definition of shape over which color pigment cells populate during maturation.

Homozygous Purple Body (*Pb/Pb*) alters all orange spots found in the body and in finnage to “pinkish-purple”, though modification may not be so readily visible in regions of red solid color. It therefore should be considered a “full body” modifier. Homozygous Pb can also produce a purple guppy phenotype. Homozygous Pb removes all visible yellow color pigment over white leucophores, but not Mg in body and finnage. This in turn, produces a dramatic increase in the visibility of wild-type violet-blue iridophores. The number of melanophores does not appear to drastically increase in any given individual as compared to homozygous Pb, but the size of the melanophores themselves was greater.

Heterozygous Pb exhibits partial reduction in collected xanthophores, and homozygous Pb has a near complete removal of collected and clustered xanthophores. However, yellow color cell populations consisting of isolated “wild-type” single cell xanthophores remain intact.

Further descriptions of Guppy Traits are available for download in: Bias and Groenewegen (2016, *with periodic updates) Poecilia reticulata:* Domestic Breeder Trait Matrix Reference Guide. https://www.academia.edu/29928596/Poecilia_reticulata_Domestic_Breeder_Trait_Matrix_Reference_Guide (last checked 1.21.2017).

### Phenotypic expression of Pb and non-Pb modification in Grey (*wild-type*)

**Fig 4.**
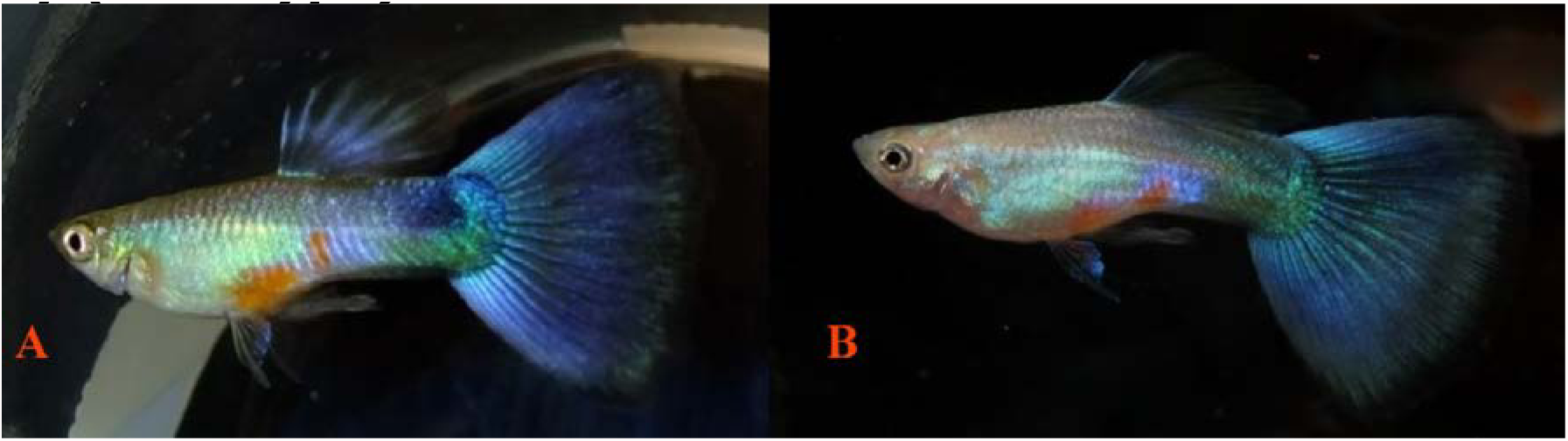
**(A)** Purple Delta (*Pb/pb*) males. Results of a homozygous Green (*pb/pb*) male x homozygous Purple (*Pb/Pb*) female breeding. **(B)** Homozygous Purple (*Pb/Pb*) male x homozygous Green (*pb/pb*) female breeding. This type male will express as either blue or purple depending upon the angle of light.

**Fig 5.**
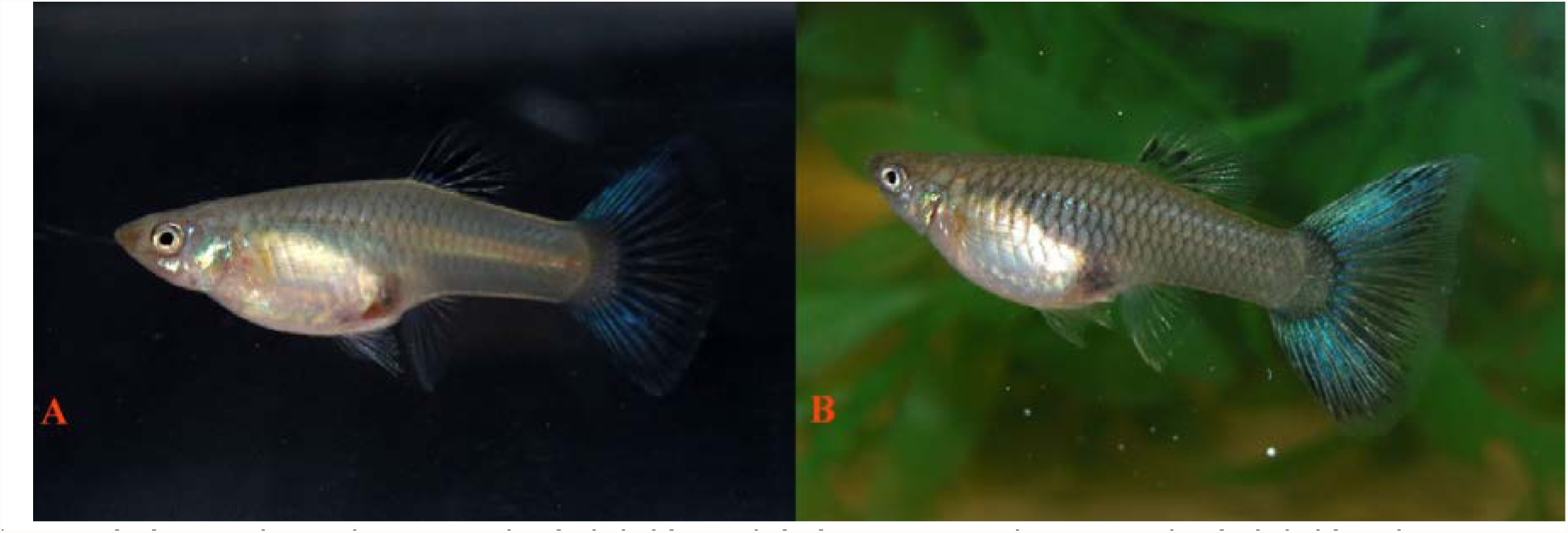
**(A)** Purple Delta Female (*Pb/Pb*) and **(B)** Green Delta Female (*pb/pb*), photos courtesy of Bryan Chin. Note the dark violet-blue color of the caudal fin with Pb, green reduced through xanthophore removal.

**Fig 6.**
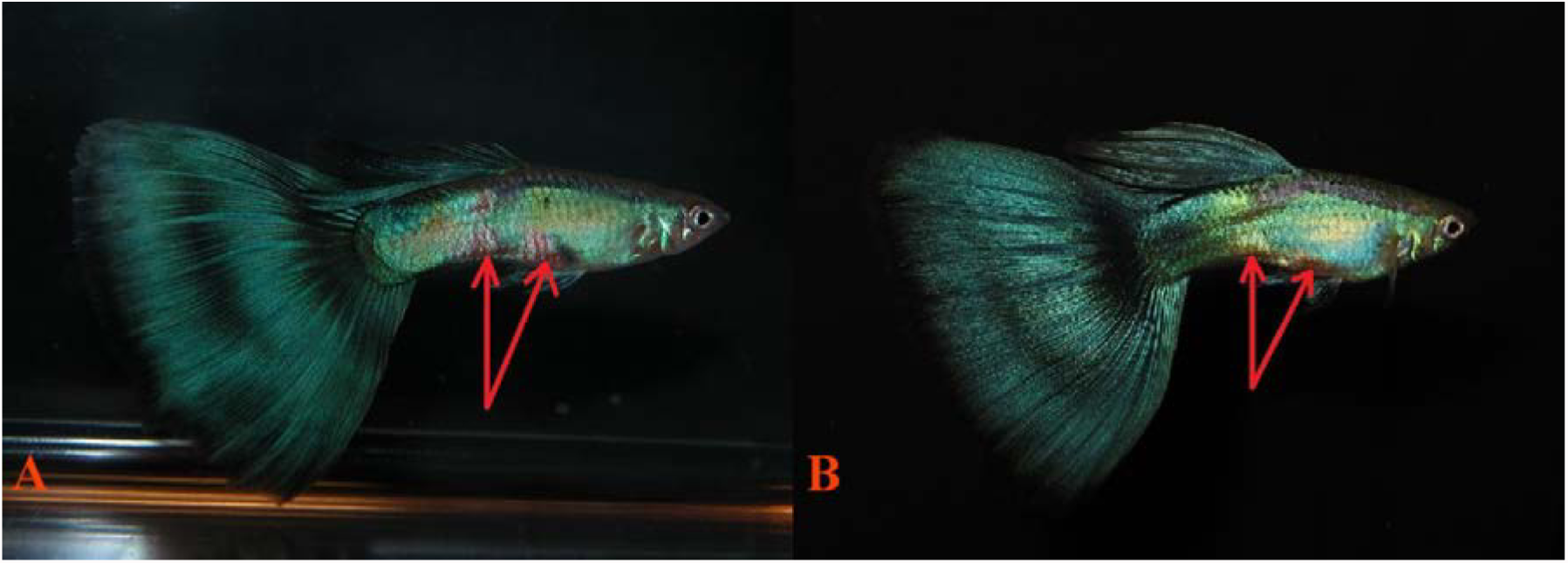
**(A)** Green Delta expressing Pb modified ornaments (*Pb/pb*). **(B)** Green Delta (*pb/pb*), photos courtesy of Bryan Chin. Note the deepening of the orange body spots to pinkish-purple with Pb through xanthophore removal (arrows).

**Fig 7.**
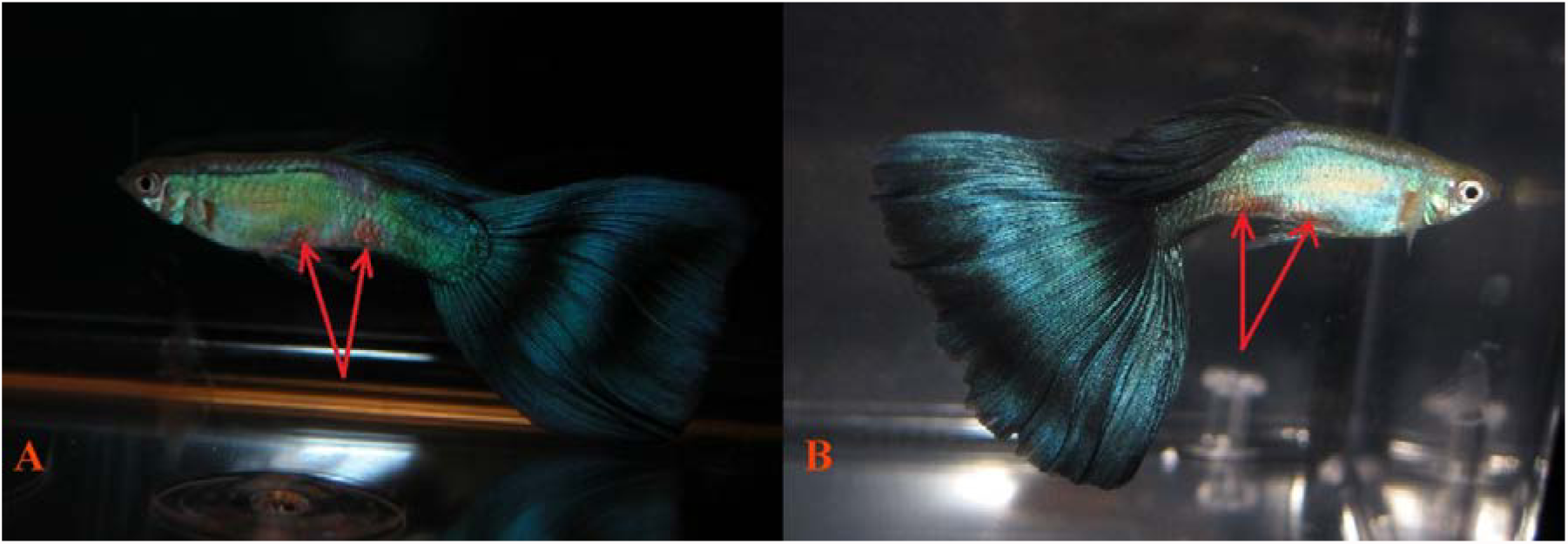
**(A)** Blue Delta expressing Pb modified ornaments (*Pb/pb*). **(B)** Blue Delta (*pb/pb*), photos courtesy of Bryan Chin. Note the deepening of the orange body spot to pinkish-purple with Pb through xanthophore removal (arrows).

**Fig 8.**
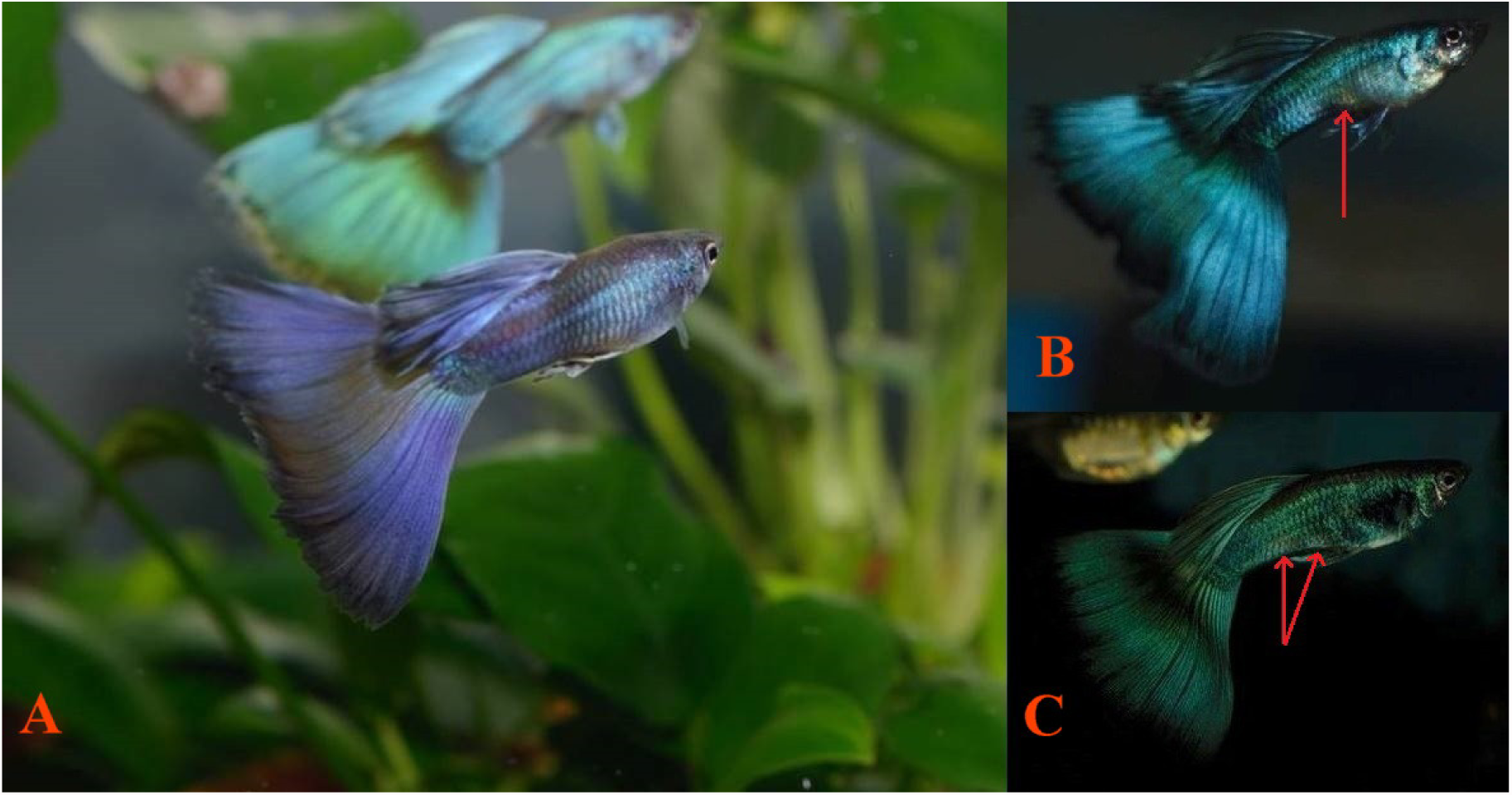
**(A)** Purple Moscow (*Mw + MBAG*) Delta expressing Pb modified ornaments (*Pb/Pb*). **(B)** Blue-Green Moscow (*Mw + MBAG*) Delta expressing Pb modified ornaments (*Pb/pb*). Note violet-blue iridophore based pattern and reduction of xanthophores in heterozygous Pb condition. Unmodified anterior orange body spot partially masked by MBAG in peduncle (arrows). **(C)** Green Moscow (*Mw + MBAG*) Delta (*pb/pb*). Note absence of Pb effects. Unmodified anterior and posterior orange body spots partially masked in peduncle by MBAG (arrows), photos courtesy of Igor Dusanic.

**Fig 9.**
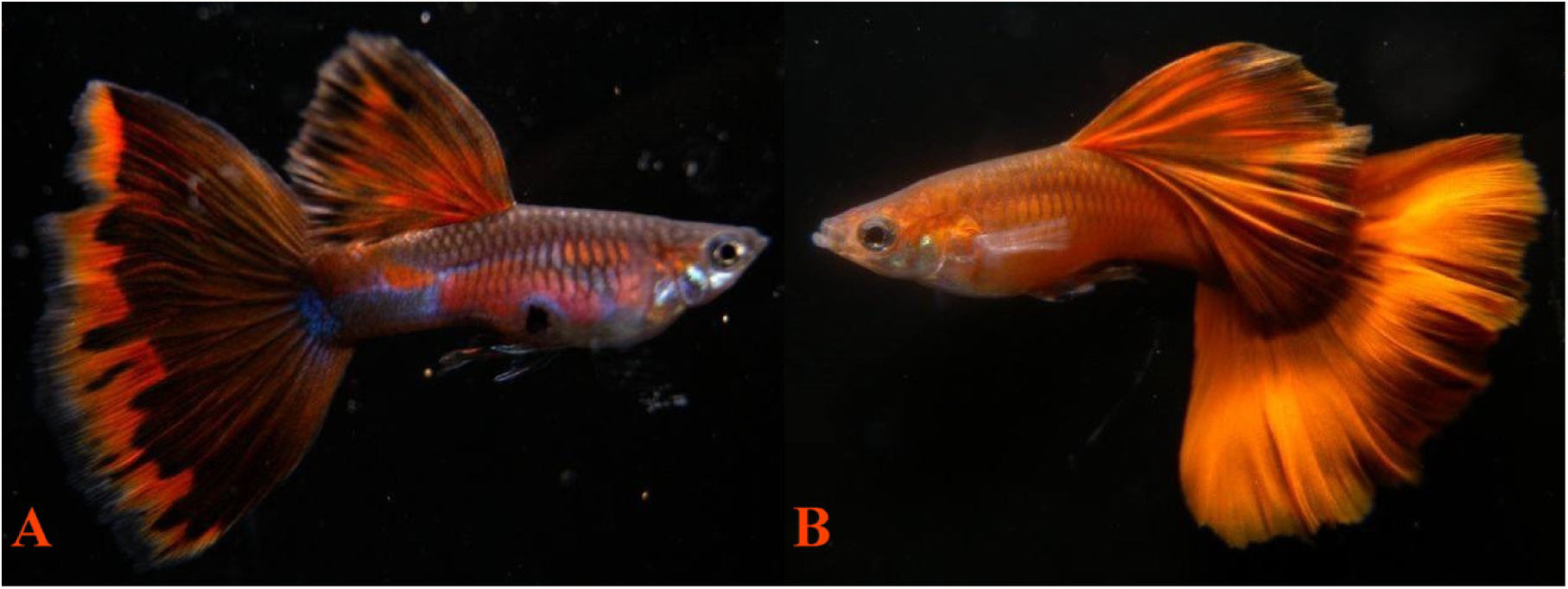
**(A)** Red Delta expressing Pb modified ornaments (*Pb/pb*). **(B)** Red Delta (*pb/pb*), photos courtesy of 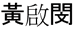 Kiddo Huang. Note how Pb darkens deep orange to dark red through xanthophore removal, and modifies body spots from orange to pinkish-purple with increase violet-blue iridophore expression.

**Fig 10.**
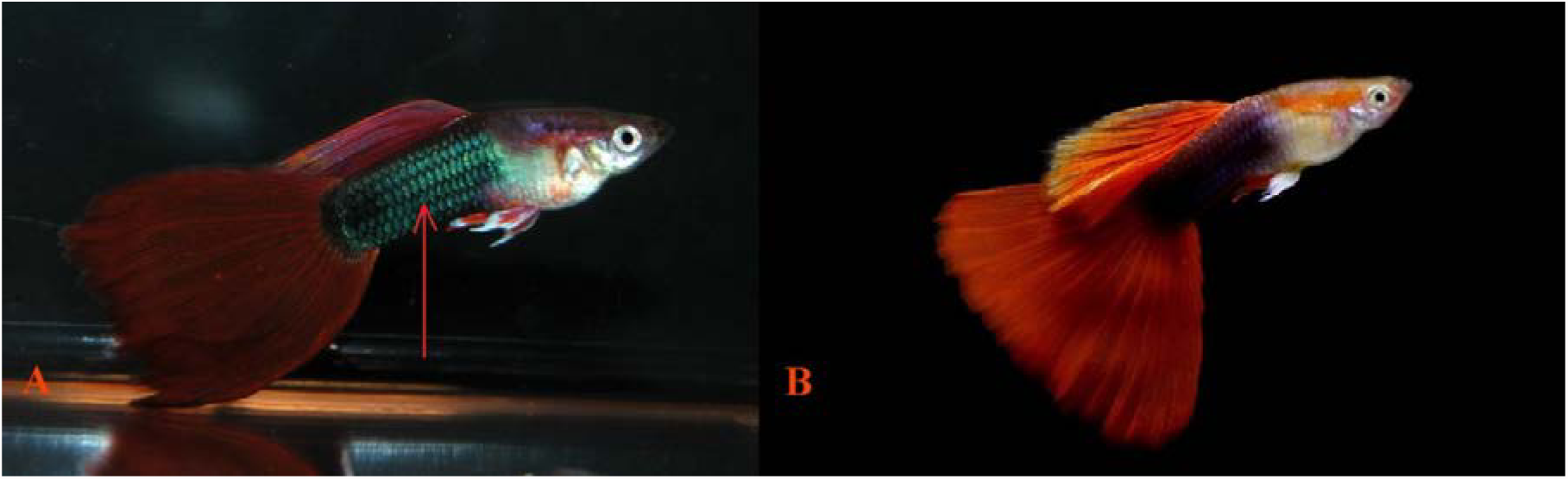
**(A)** Half Black (*NiII*) Red Delta expressing Pb modified ornaments (*Pb/pb*). **(B)** Half Black (*NiII*) Red Delta (*pb/pb*), photos courtesy of Bryan Chin and Cheng-Hsien Yang. Note the darker red replacing lighter orange-red by Pb through xanthophore removal. Peduncle and topline expresses increased iridophores in Pb, and dorsal shows reduction of xanthophores revealing violet-blue iridophores (arrow).

**Fig 11.**
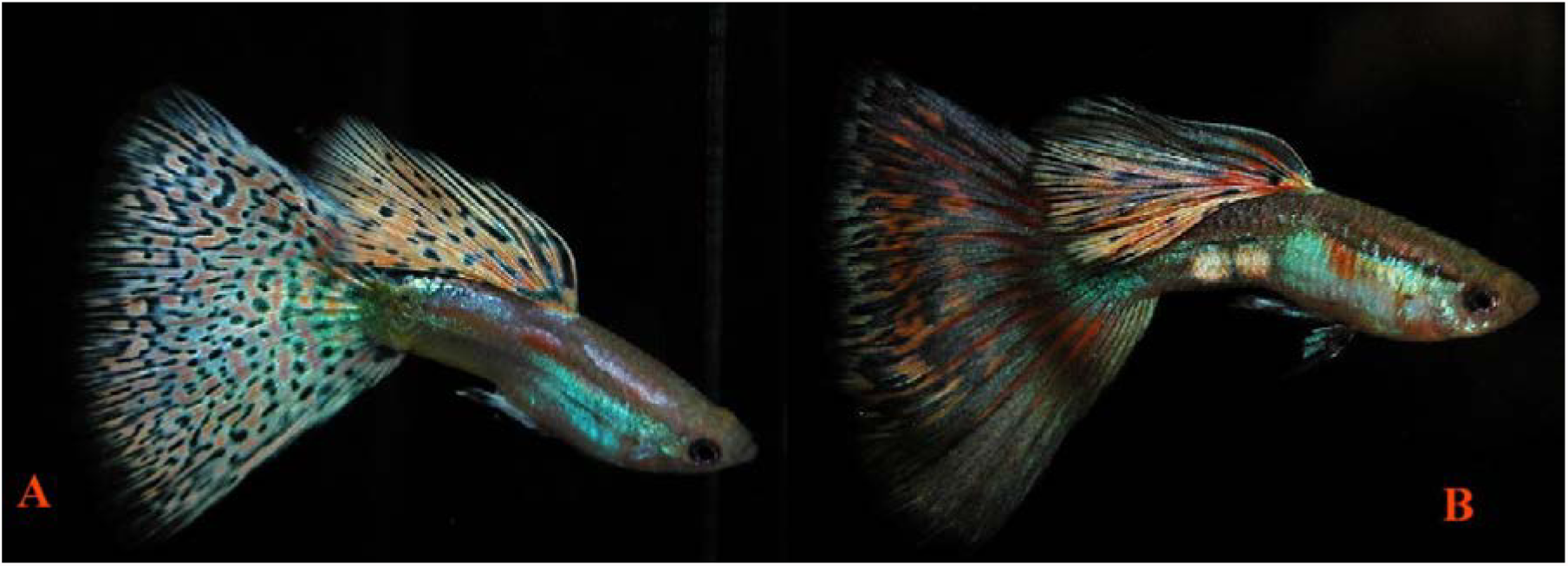
**(A)** Red Multi-Color (*Pb/pb*) Delta expressing Pb modified ornaments. Note the change of orange body spots to pinkish-purple from xanthophore removal, while darker red is unaffected with Pb. **(B)** Red Multi-Color Delta (*pb/pb*), photos courtesy of Bryan Chin.

**Fig 12.**
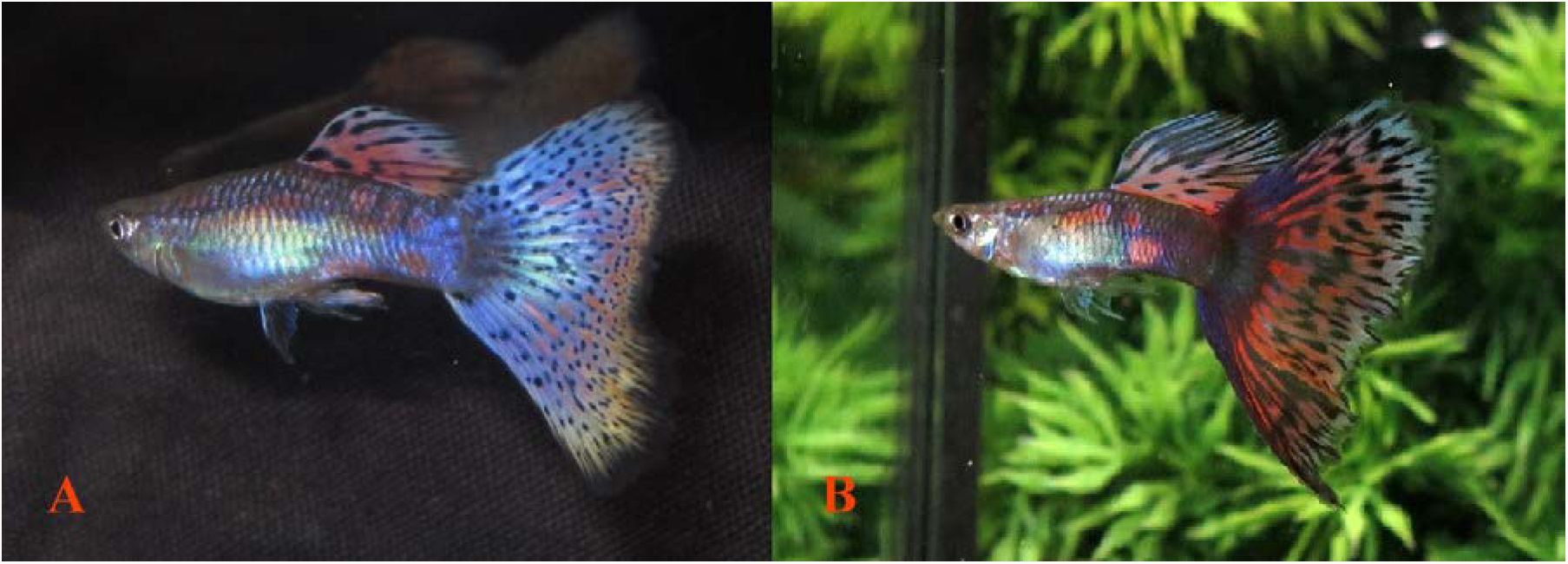
**(A)** Purple Multi-Color (*Pb/Pb*). **(B)** Red Multi-Color Deltas expressing Pb modified ornaments (*Pb/Pb*), photos courtesy of Bryan Chin. Note the erythrophores show partial Pb modification in both males’ finnage and the violet coloration in the fish to the left. Genes other than Pb are responsible for some of these differences. Some orange spots are modified to pinkish-purple. Pb reduces sex-linked xanthophores in dorsal and caudal, revealing white leucophores (*Le*) in finnage. There is a slight increase in violet-blue structural color in body.

**Fig 13.**
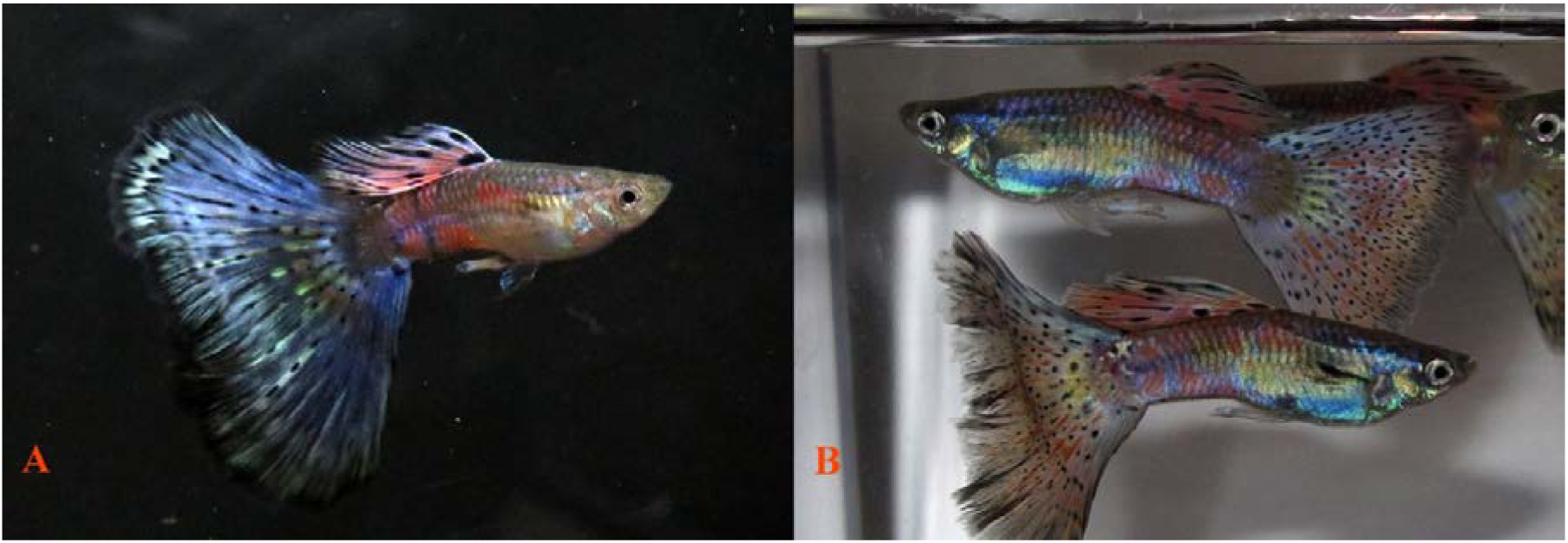
**(A)** Purple Multi-Color Delta (*Pb/Pb*). **(B)** Red Multi-Color Deltas expressing Pb modified ornaments (*Pb/Pb*), photos courtesy of Bryan Chin. Again, additional genes are involved here. Orange spots are modified to pinkish-purple. Pb reduces sex-linked xanthophores in dorsal and caudal, revealing white leucophores (*Le*) in finnage. There is a slight increase in violet-blue structural color in body.

**Fig 14.**
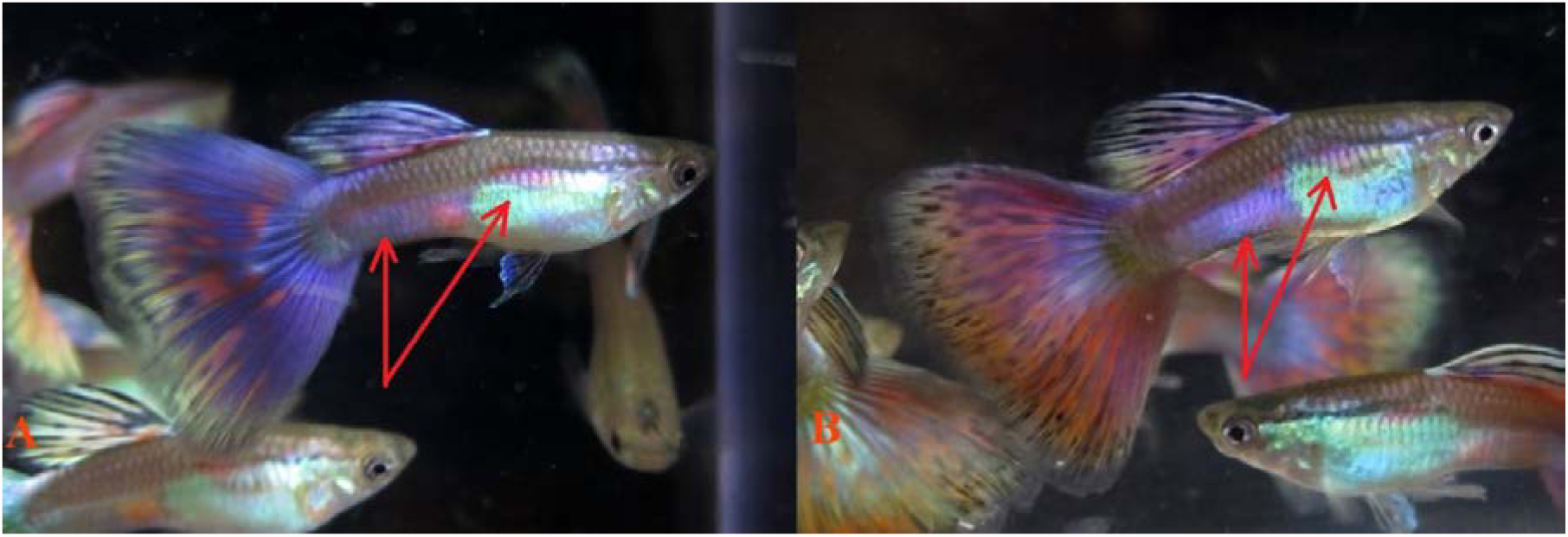
**(A)** Purple Multi-Color (*Pb/Pb*). **(B)** Red Multi-Color Deltas expressing Pb modified ornaments (*Pb/Pb*), photos courtesy of Bryan Chin. Orange spots are modified to pinkish-purple, and increased violet-blue iridophores (arrows). Some anterior Metal Gold (Mg) remains over blue iridophores and in the VEG peduncle spot.

**Fig 15.**
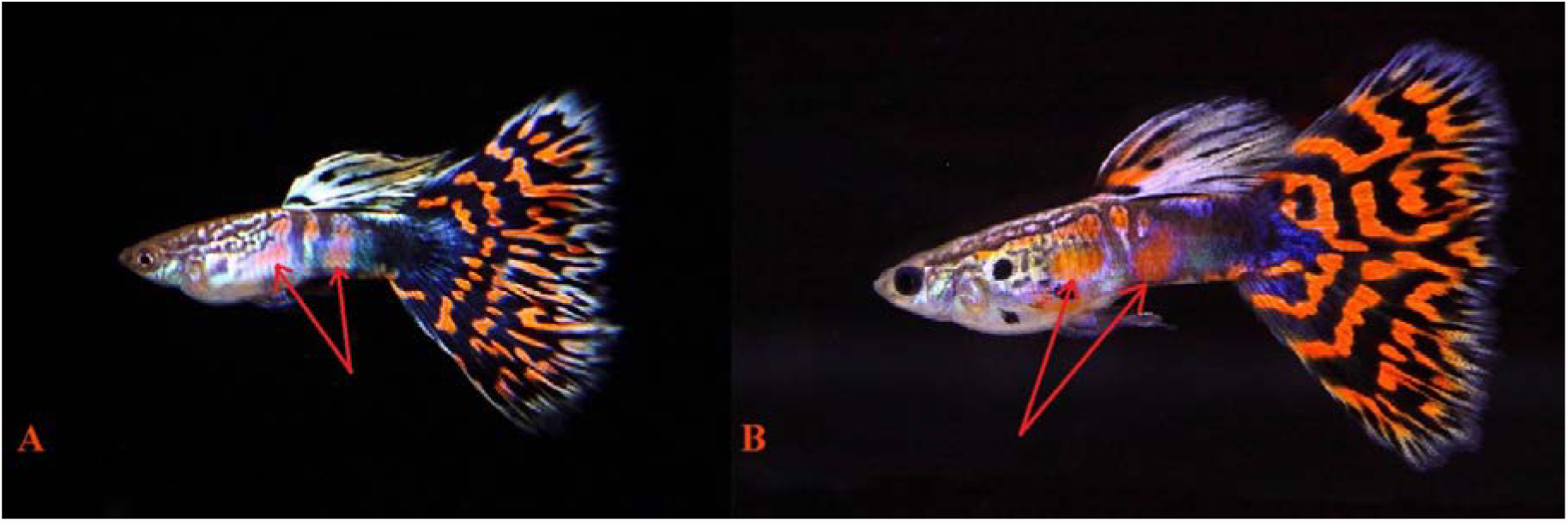
**(A)** Red Mosaic Delta (*Pb/Pb or Pb/pb*) expressing Pb modified ornaments. **(B)** Red Mosaic Delta (*pb/pb*), photos courtesy of Kevin and Karen Yang. Pb converts lighter carotenoid orange to pinkish-purple on the body (arrows), but the effect on the darker pteridine red in caudal fin is less pronounced. Pb reduces sex-linked xanthophores in dorsal and caudal, revealing white leucophores (*Le*) in finnage. There is a slight increase in violet-blue structural color in body.

**Fig 16.**
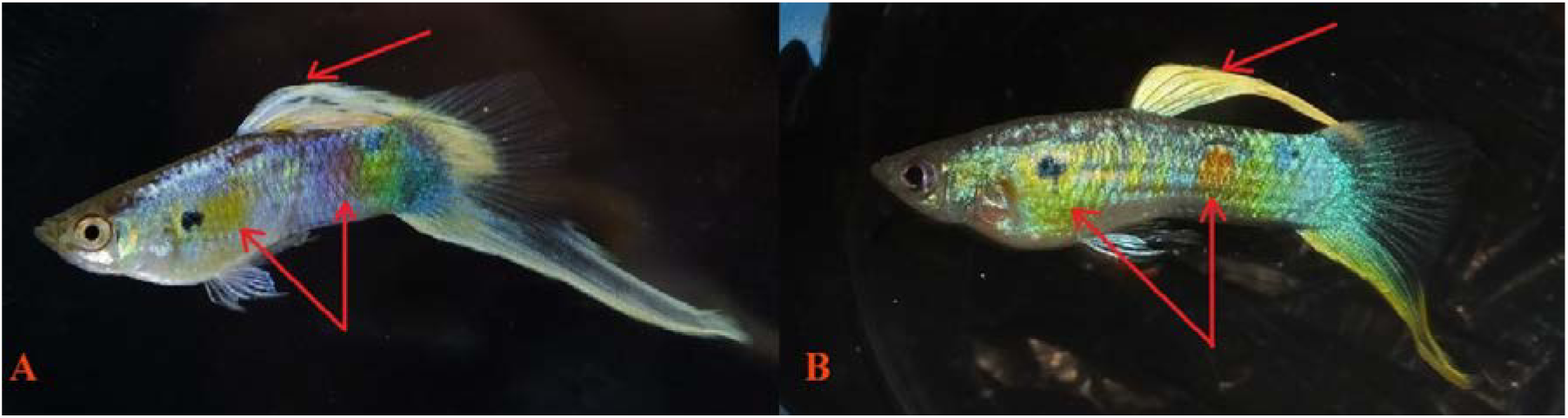
**(A)** Vienna Lowersword expressing Pb modified ornaments (*Pb/Pb*). **(B)** Vienna Lowersword (*pb/pb*). Green coloration is the product of yellow xanthophores over blue iridophores. Note that when Pb reduces the yellow, it reduces green as well. Posterior orange spots are modified to pinkish-purple (arrows). There is increased expression of violet-blue iridophores while collected sex-linked xanthophores are removed and clustered Metal Gold (*Mg*) are only reduced by Pb (arrow).

**Fig 17.**
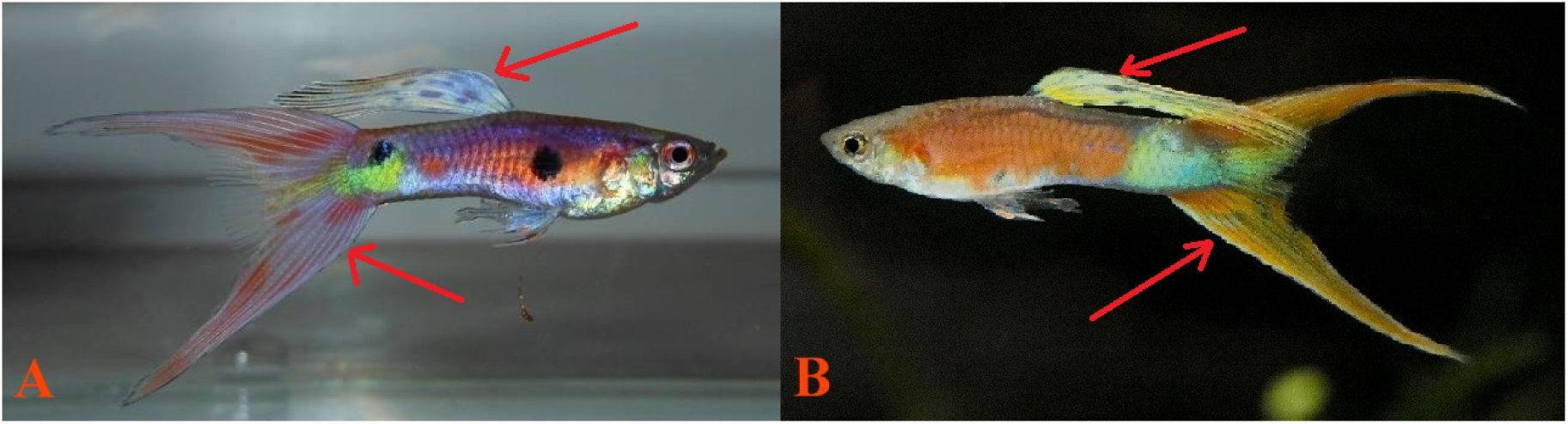
**(A):** Coral Red (*Co*) Doublesword expressing Pb modified ornaments (*Pb/Pb or Pb/pb*). **(B)** Coral Red (*Co*) Doublesword (*pb/pb*), photos courtesy of Krisztián Medveczki and Gary Lee. Note how Pb changes orange to deep red. Pb removes sex-linked xanthophores in the dorsal and caudal, leaving only white leucophores (*Le*), as no sex-linked erythrophores are present. Some orange spots are modified to pinkish-purple, and there is proliferation of iridophores in the body. Sex-linked collected yellow pigment is removed from both caudal and dorsal (arrow). There is a heavy increase in violet-blue structural color in body, but not finnage, as Pb modification of erythrophores is minimal.

**Fig 18.**
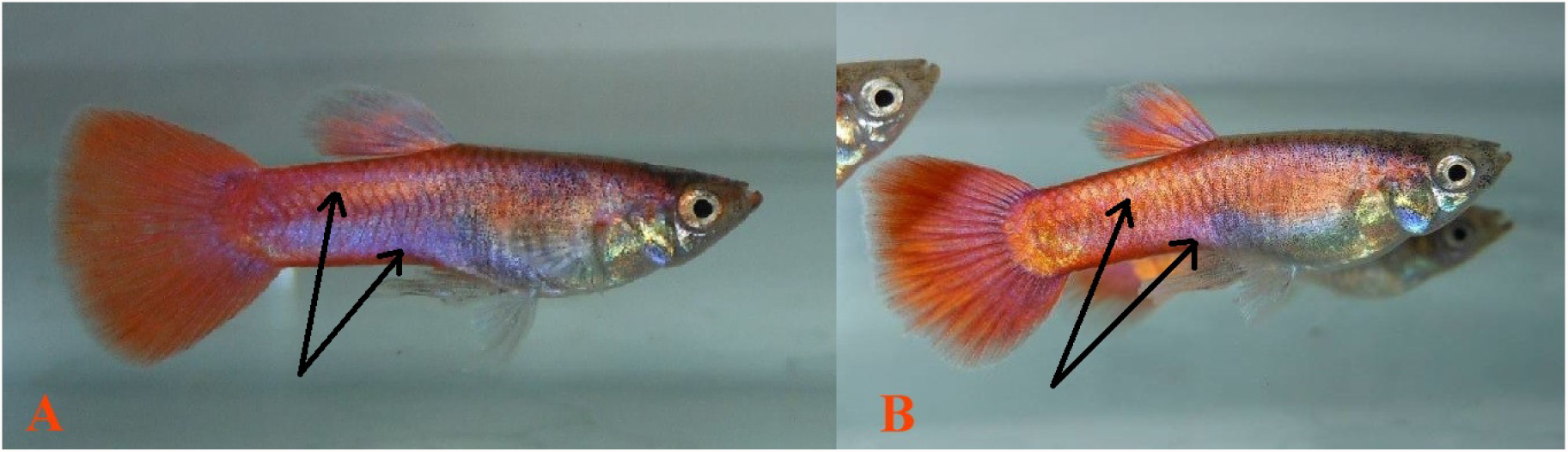
**(A)** Magenta (*M*) Delta expressing Pb modified ornaments (*Pb/pb*). **(B)** Magenta (*M*) Delta (*pb/pb*), photos courtesy of Krisztián Medveczki. Note how Pb converts orange to red by reduction of xanthophores (arrows) and deepens and expands the violet-blue iridophore coloration. Orange spots are modified to pinkish-purple, and increased expression of violet-blue iridophores by xanthophore reduction (arrows).

**Fig 19.**
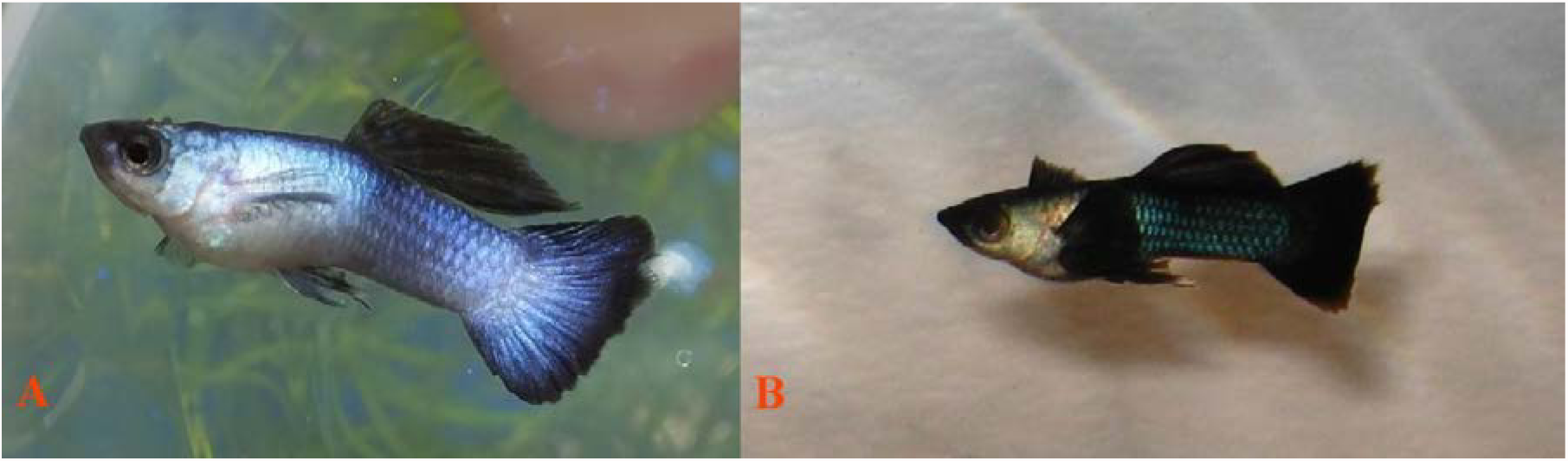
**(A)** Purple Panda Moscow (*Mw + pp*) Roundtail expressing Pb modified ornaments (*Pb/Pb*). **(B)** Panda Moscow (*Mw + pp*) Roundtail (*pb/pb*). Green is replaced by purple with Pb, by removal of xanthophores with increased expression of violet-blue iridophores.

### Phenotypic expression of Pb and non-Pb modification in Albino (*aa*)

**Fig 20.**
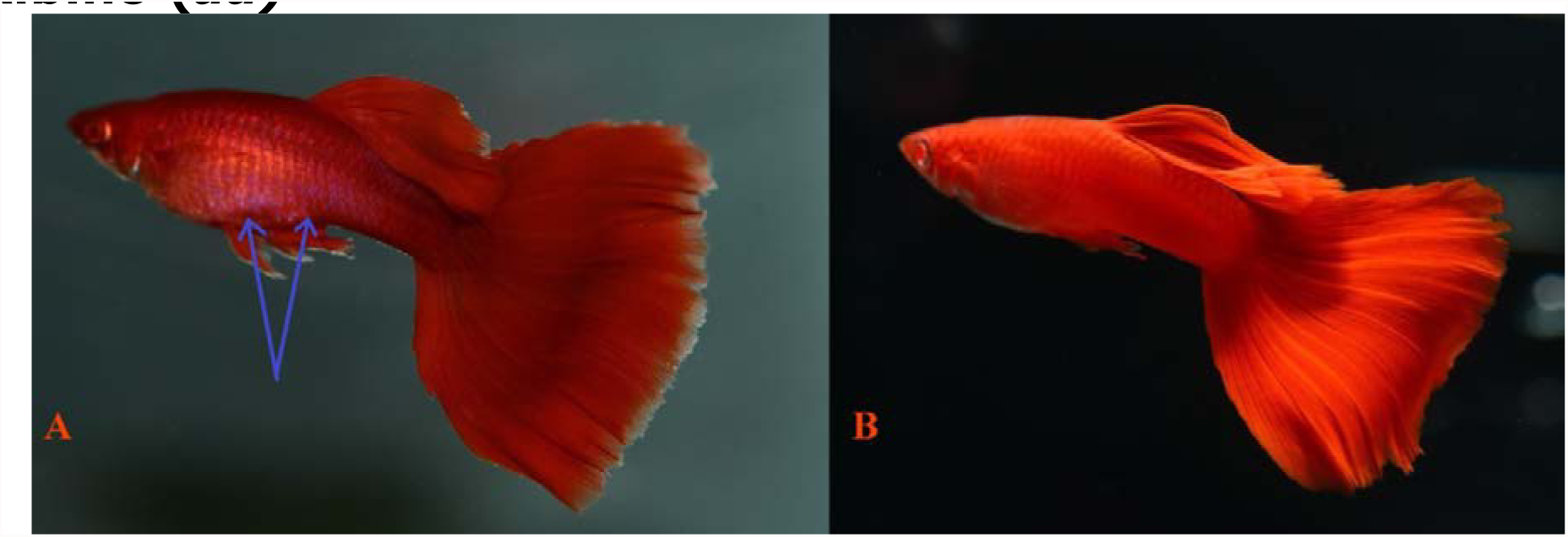
**(A)** Full Red Albino (aa) Delta expressing Pb modified ornaments (*Pb/pb*). **(B)** Full Red Albino (aa) Delta (*pb/pb*), photos courtesy of 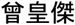 Tseng Huang Chieh. Melanophores are removed by Albino. **(A)** Orange red is changed to dark red by Pb through xanthophore removal, with proliferation of violet-blue iridophores visibly modified to pinkish-purple (arrows).

**Fig 21.**
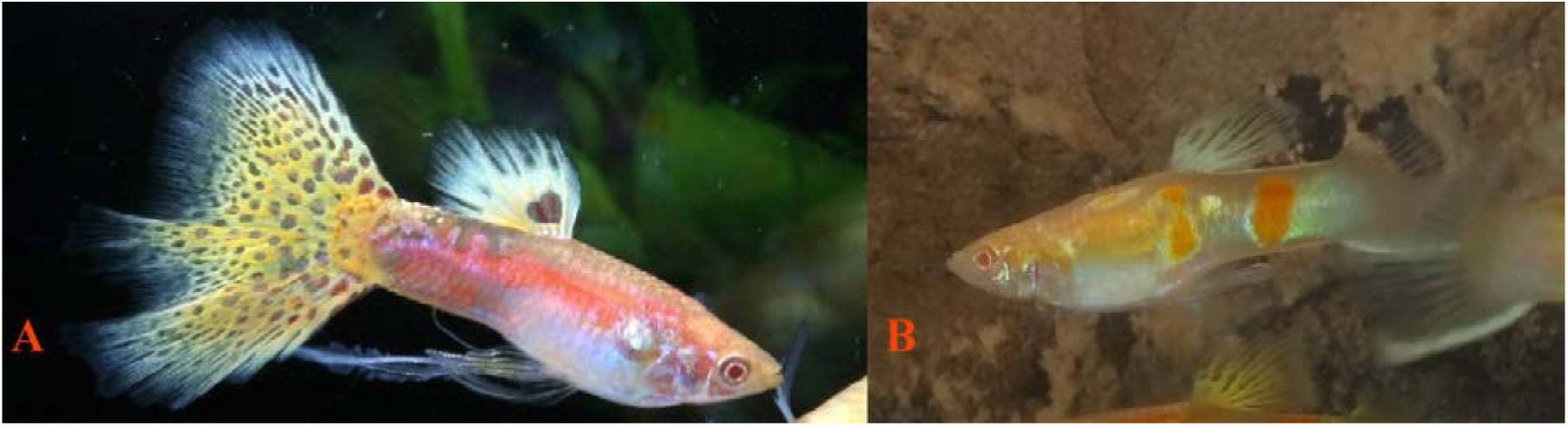
**(A)** Albino (aa) Delta expressing Pb modified ornaments (*Pb/Pb or Pb/pb*), photo courtesy of Benson Liu. **(B)** Albino (*aa*) Lowersword (*pb/pb*). Melanophores are removed by Albino, though dendritic pattern remains. Posterior orange spots are modified to pinkish-purple, with increased expression of violet-blue iridophores.

### Phenotypic expression of Pb and non-Pb modification in Blond (*bb*)

**Fig 22.**
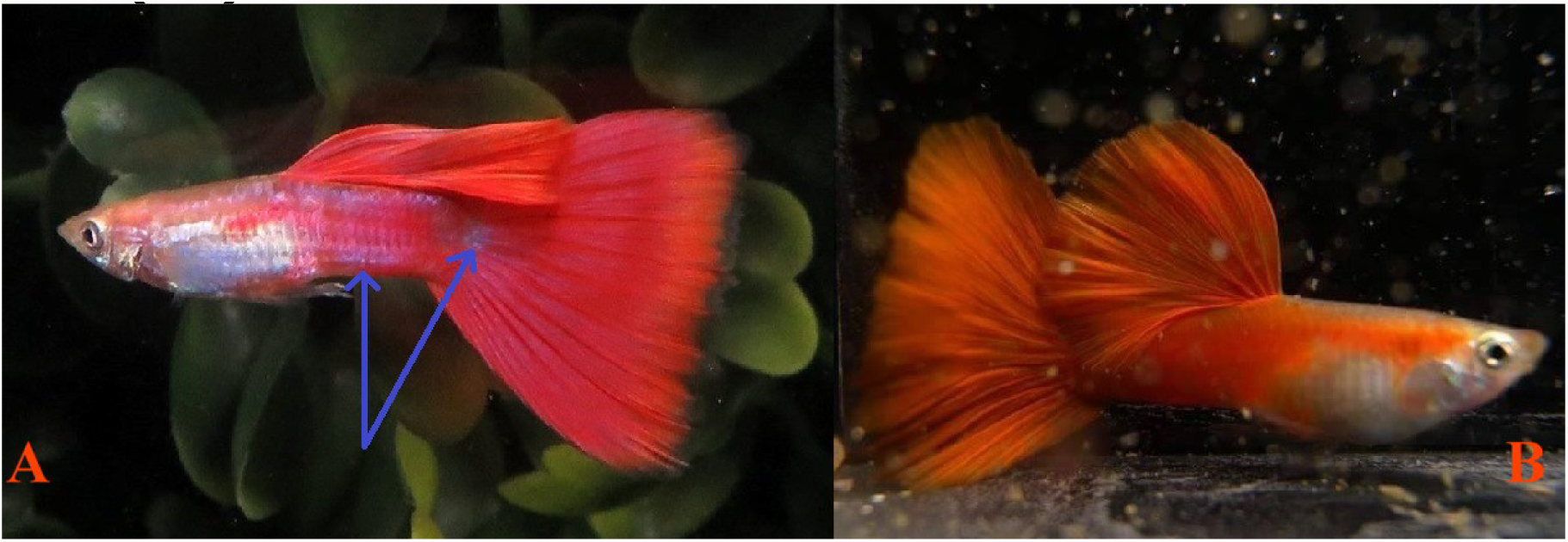
**(A)** Blond (*bb*) Red Delta expressing Pb modified ornaments (*Pb/pb or Pb/Pb*). **(B)** Blond (*bb*) Red Delta (*pb/pb*), photos courtesy of Bryan Chin and 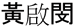 Kiddo Huang. Removal of xanthophores and a proliferation of violet-blue iridophores modifies orange to a darker red and spots are modified to pinkish-purple (arrows).

**Fig 23.**
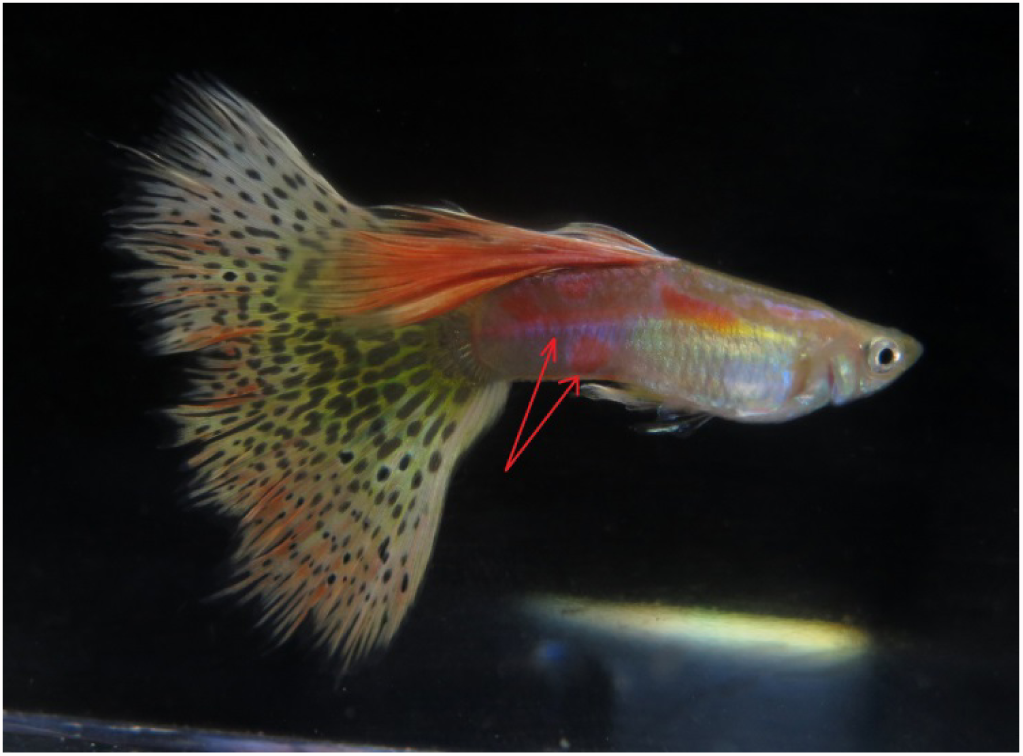
Blond (*b*) Red Multi-Color Delta expressing Pb modified ornaments (*Pb/pb*), photo courtesy of Bryan Chin. Blond reduces melanophore size. Posterior orange is converted to pinkish-purple by Pb (arrows), with increased expression of violet-blue iridophore structural color. Pb reduces sex-linked xanthophores in dorsal and caudal, revealing white leucophores (**Le**) in finnage.

**Fig 24.**
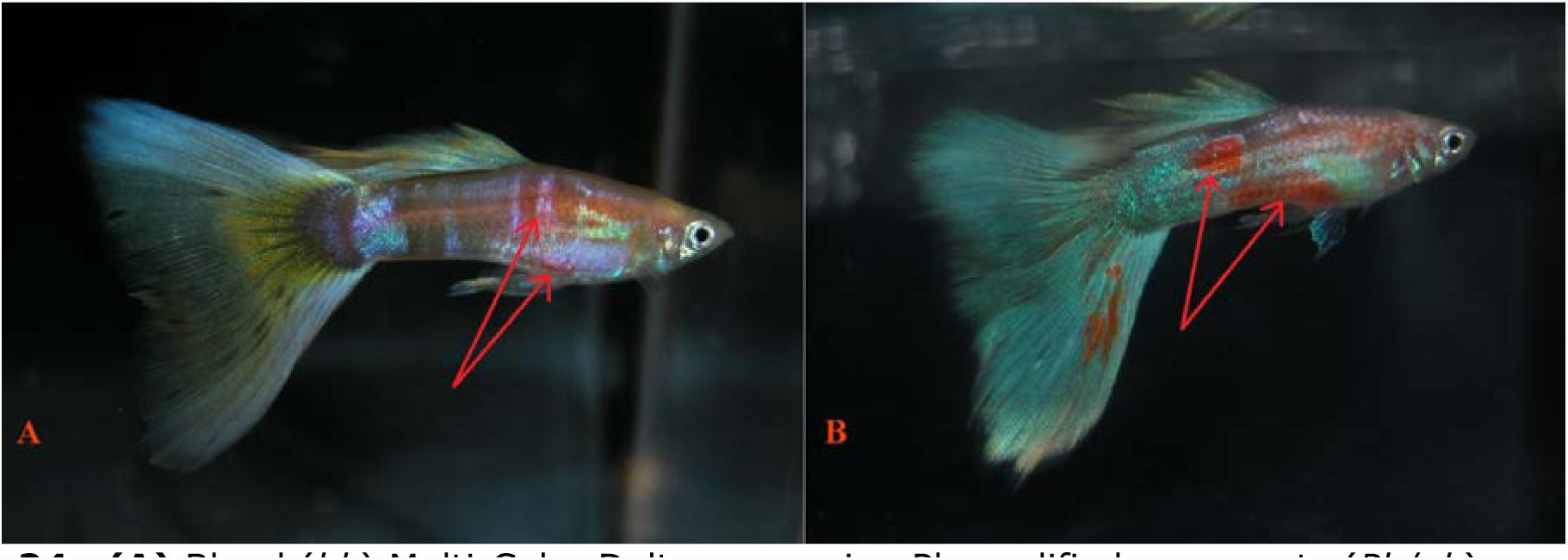
**(A)** Blond (*bb*) Multi-Color Delta expressing Pb modified ornaments (*Pb/pb*). Heterozygous Pb converts some orange spots over Zebrinus (Ze) to pinkish-purple (arrows) with increased purple violet iridophores, some anterior orange remains. Pb reduces sex-linked xanthophores in dorsal and caudal fins, revealing white leucophores (**Le**) in finnage. There is a slight increase in violet-blue structural color in body. **(B)** Blond (*bb*) Multi-Color Delta (*pb/pb*), photos courtesy of Bryan Chin. Blond reduces melanophore size. There is no modification to orange spotting ornaments (arrows) comprised of xantho-erythrophores.

**Fig 25.**
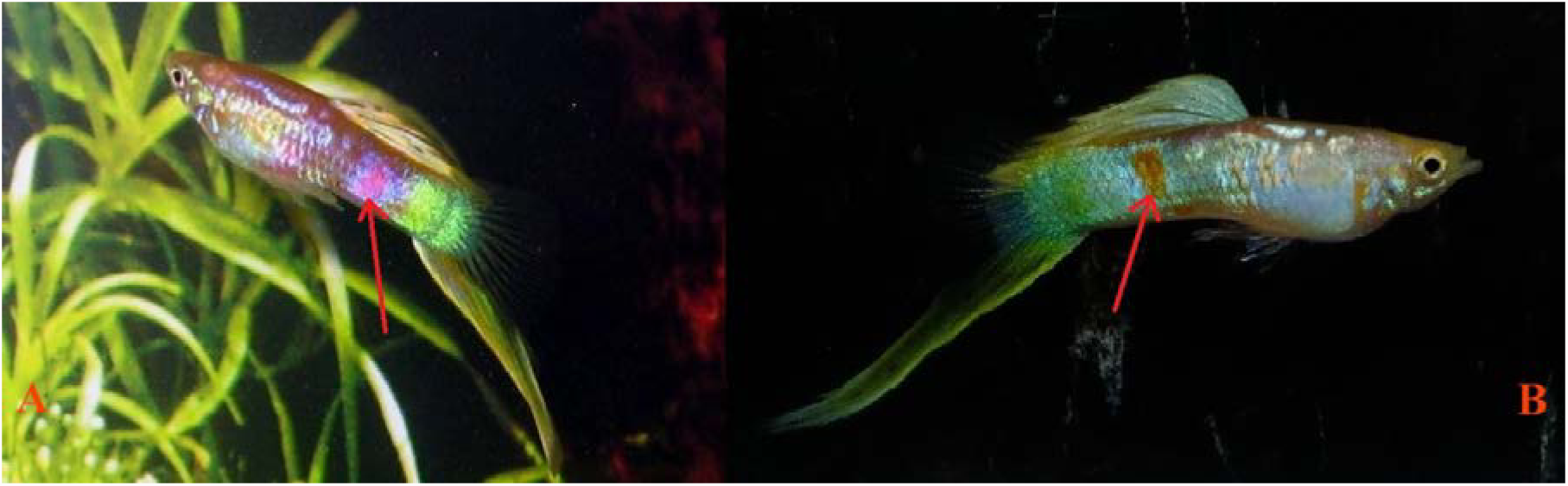
**(A)** Blond (*bb*) Vienna Lowersword expressing Pb modified ornaments (*Pb/Pb*). **(B)** Blond (*bb*) Vienna Lowersword (*pb/pb*). Blond reduces melanophore size. Orange is converted to pinkish-purple by xanthophore removal (arrows), green is almost eliminated by Pb.

**Fig 26.**
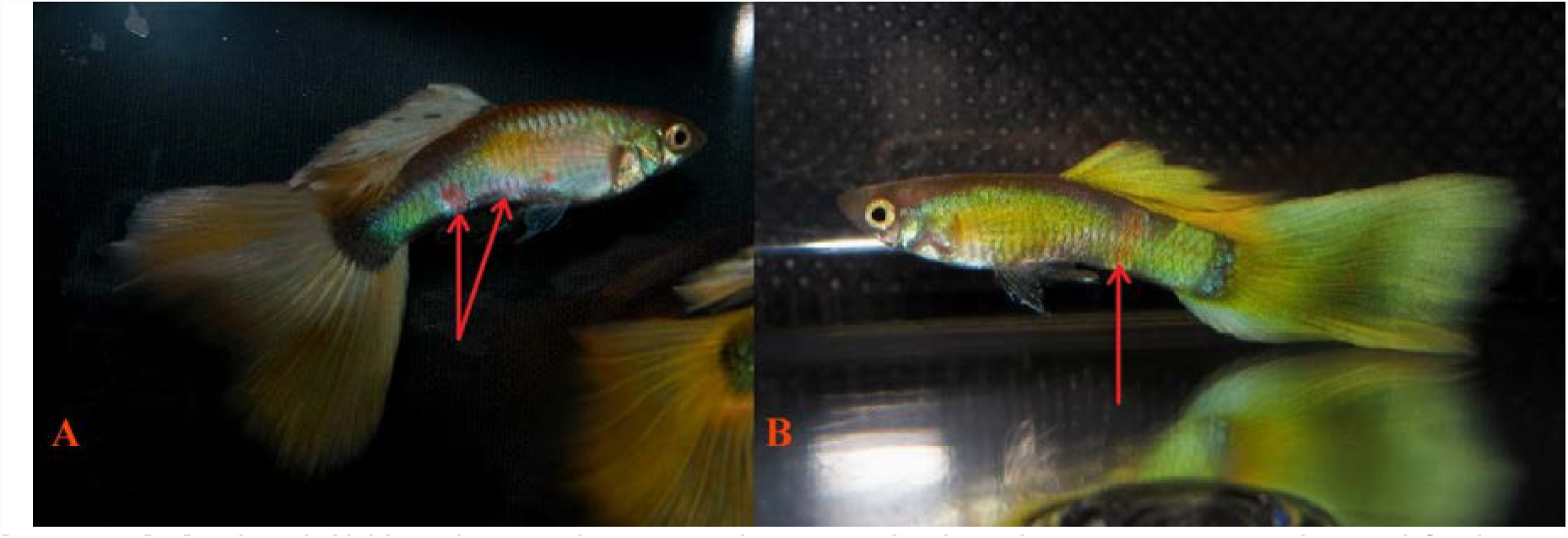
**(A)** Blond (*bb*) Schimmelpennig Platinum (*Sc*) Delta expressing Pb modified ornaments (*Pb/Pb*). **(B)** Blond (*bb*) Schimmelpennig Platinum (*Sc*) Delta (*pb/pb*), photos courtesy of Bryan Chin. Blond reduces melanophore size. Orange is converted to pinkish-purple by xanthophore removal (arrows), green is eliminated by Pb. Pb reduces sex-linked xanthophores in dorsal and caudal fins, revealing white leucophores (**Le**) in finnage. Metal Gold (*Mg*) remains in body and finnage. There is a slight increase in violet-blue structural color in body.

**Fig 27.**
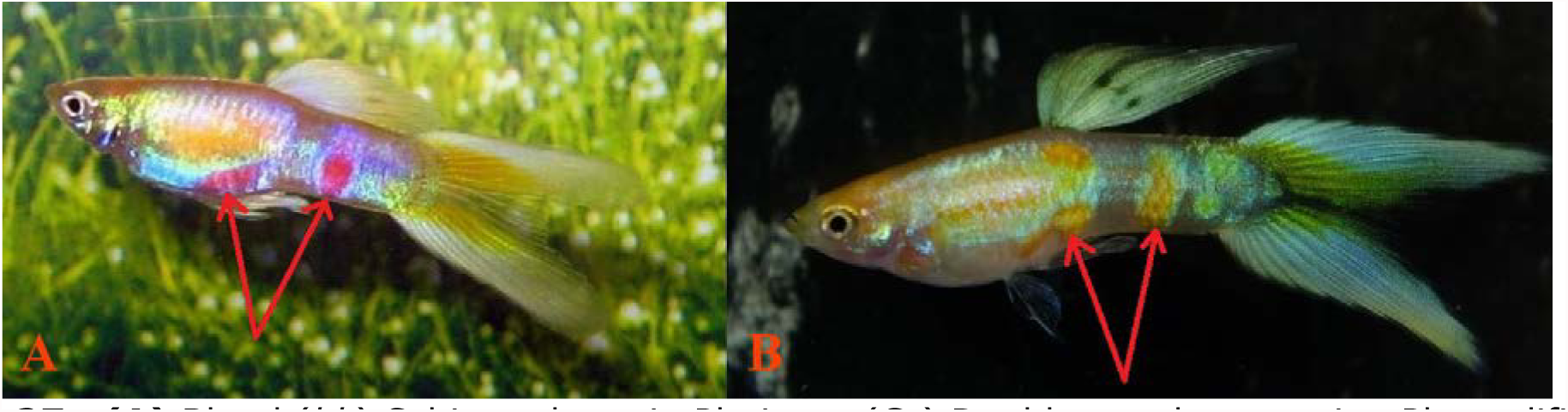
**(A)** Blond (*bb*) Schimmelpennig Platinum (*Sc*) Doublesword expressing Pb modified ornaments (*Pb/Pb*). **(B)** Blond (*bb*) Schimmelpennig Platinum (*Sc*) Doublesword (*pb/pb*). Blond reduces melanophore size. Orange is converted to pinkish-purple by xanthophore removal (arrows), green is eliminated by Pb. Note that the large platinum shoulder area, comprised of clustered Metal Gold (**Mg**) xanthophores, is still present in homozygous Pb. Pb reduces sex-linked xanthophores in dorsal and caudal, revealing white leucophores (**Le**) in finnage. There is a slight increase in violet-blue structural color in body.

### Phenotypic expression of Pb and non-Pb modification in Golden (*gg*)

**Figure 28.**
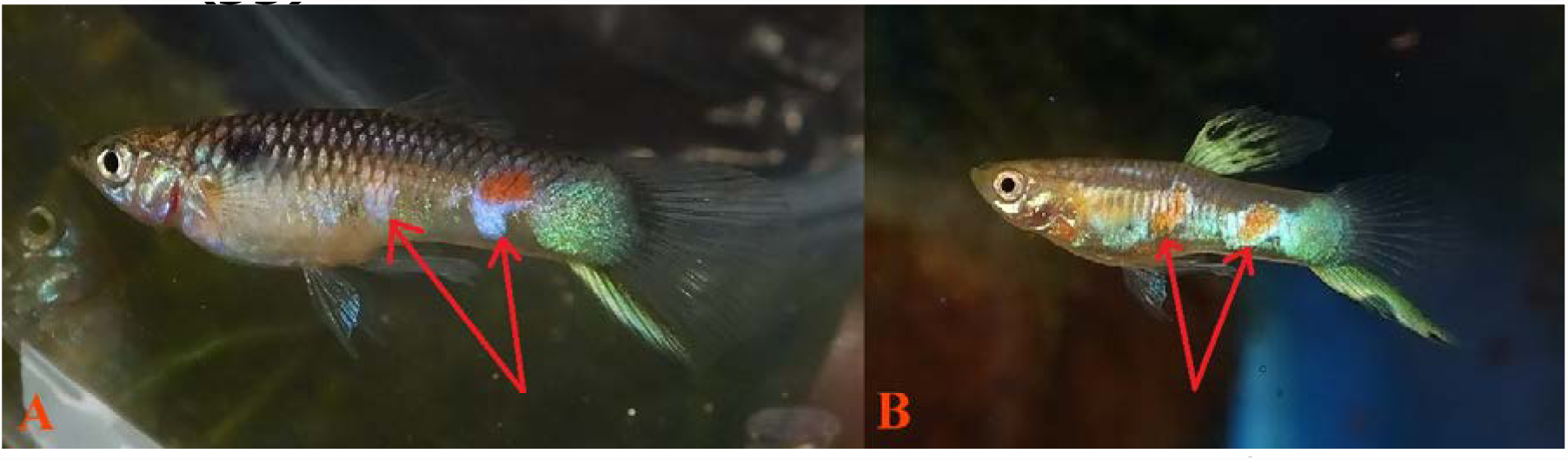
**(A)** Golden (*gg*) Vienna Lowersword expressing Pb modified ornaments (*Pb/Pb*). **(B)** Golden (*gg*) Vienna Lowersword (*pb/pb*). Melanophores are reduced and collected by Golden. Orange is converted to pinkish-purple by Pb by xanthophore removal (arrows), pale blue is deepened to violet, green is reduced. Pb reduces sex-linked xanthophores in dorsal and caudal, revealing white leucophores (**Le**) in finnage. Metal Gold (**Mg**) remains in body and finnage. Slight increase in violet-blue structural color in body.

**Fig 29.**
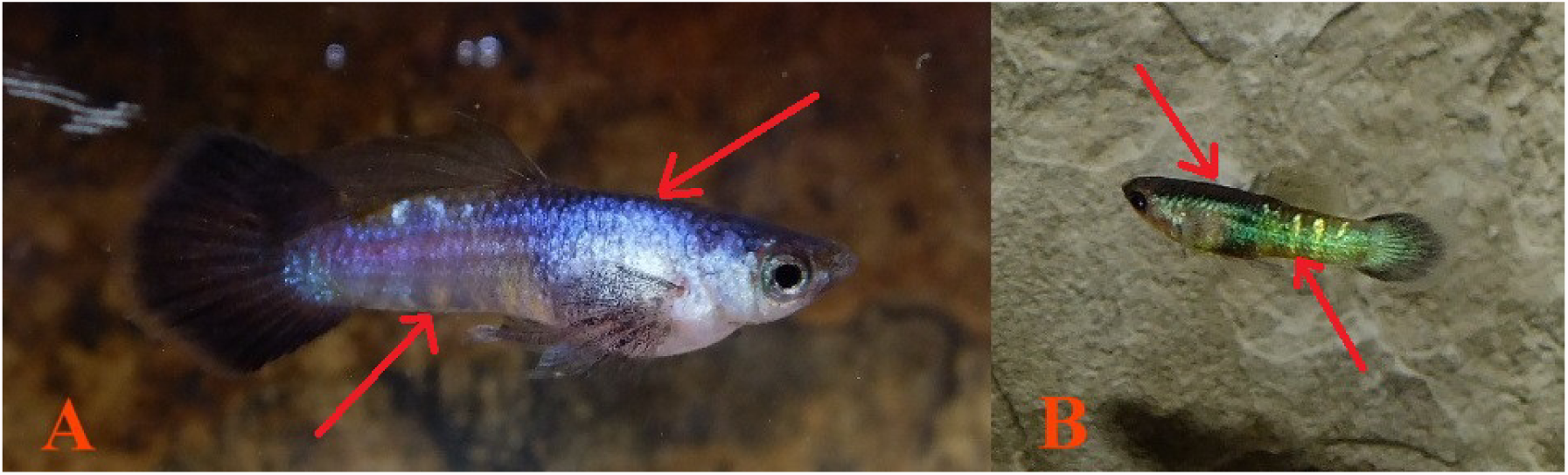
**(A)** Golden (*gg*) Purple Panda Moscow (*Mw + pp*) Roundtail expressing Pb modified ornaments (*Pb/Pb*). **(B)** Golden (*gg*) Panda Moscow (*Mw + pp*) Roundtail (*pb/pb*). Green is replaced by purple with Pb, through xanthophore removal and proliferation of violet-blue iridophores (arrow). Reduction in melanophore numbers, especially ectopic, reduces dark peduncle coloration (arrow).

### Phenotypic expression of Pb and non-Pb modification in Asian Blau (*Ab*)

**Fig 30.**
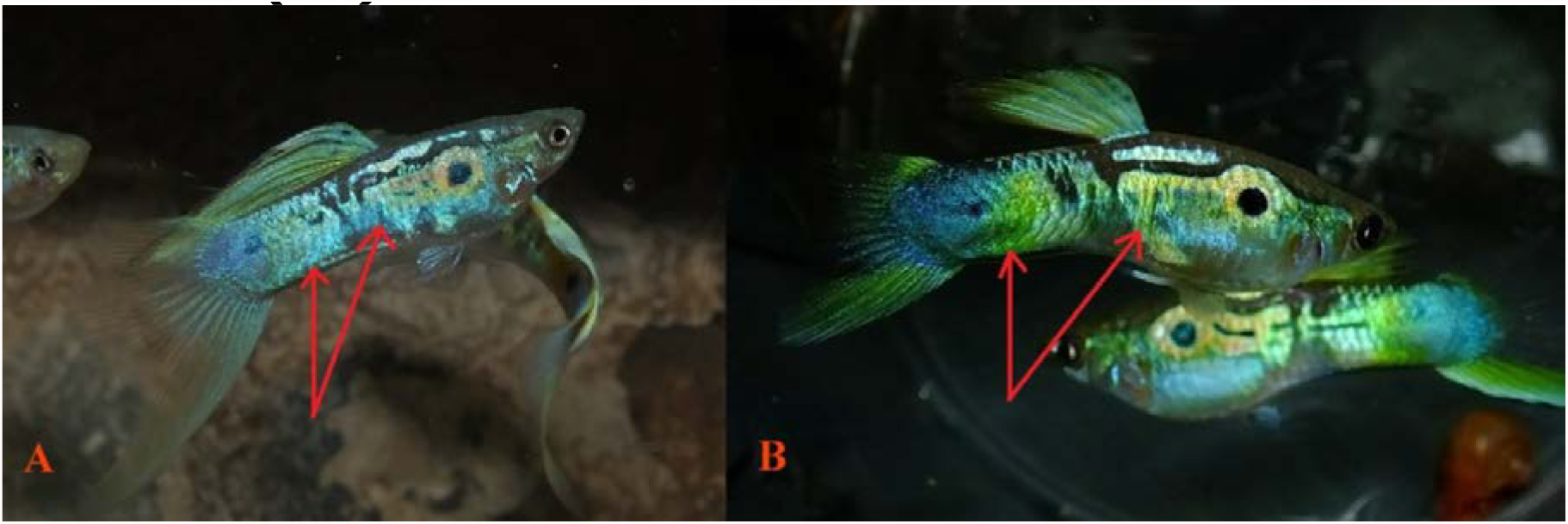
**(A)** Asian Blau (*Ab*) Vienna Lowersword expressing Pb modified ornaments (*Pb/Pb*. **(B)** Asian Blau (*Ab*) Vienna Lowerswords (*pb/pb*). Orange is converted to pinkish-purple by Pb xanthophore removal (arrows). Erythrophores are further removed by Ab revealing modified violet-blue iridophores (arrows).

**Fig 31.**
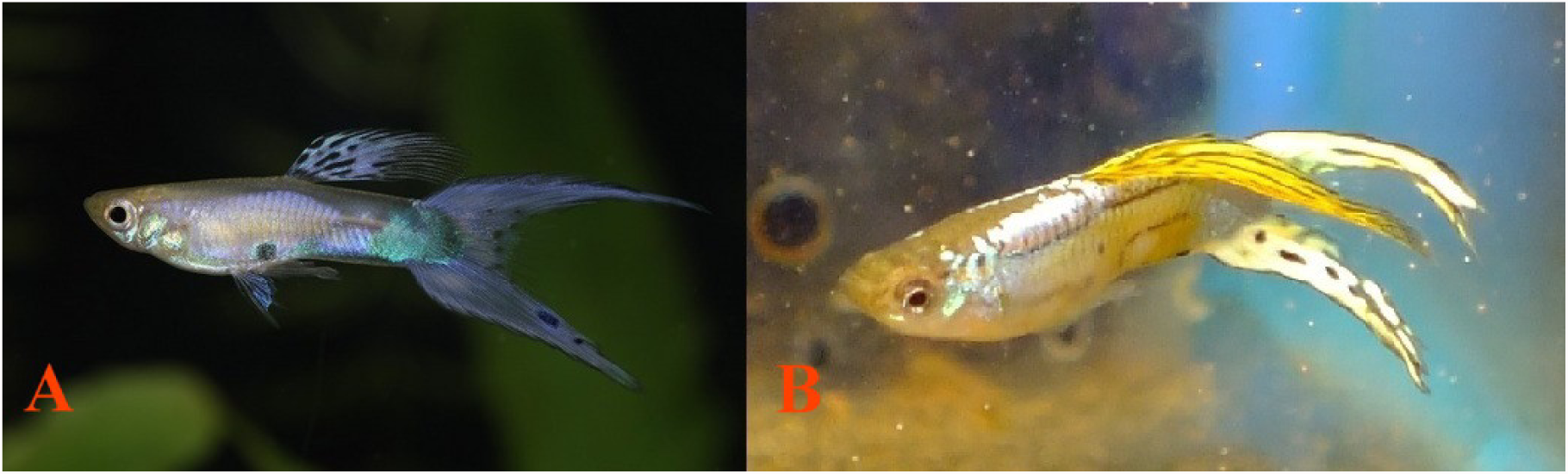
**(A)** Asian Blau (*Ab*) Ivory (*ii*); i.e. Lavender Coral Red (*Co*) Doublesword expressing Pb modified ornaments (*Pb/Pb or Pb/pb*), photo courtesy of Taketoshi Sue. Orange is converted to pinkish-purple by Pb. Asian Blau removes orange and Ivory removes yellow-orange revealing modified violet-blue iridophores and white leucophores. **(B)** Asian Blau (*Ab*) Coral Red (*Co, pb/pb*). Doublesword Orange converted to pinkish-purple by Pb, and removed by Ab revealing underlying leucophores and modified iridophores.

**Fig 32.**
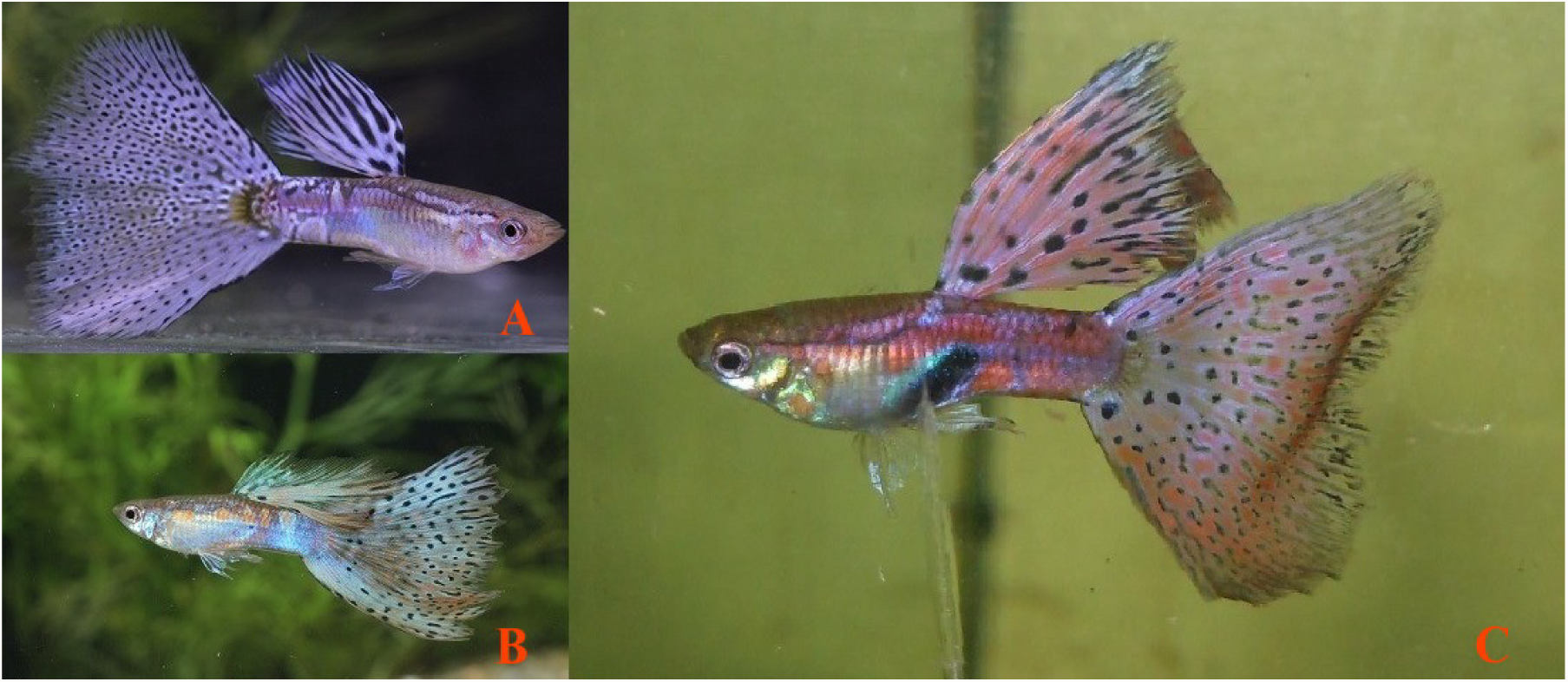
**(A)** Asian Blau (*Ab*) Ivory (*ii*); i.e. Lavender Grass Delta expressing Pb modified ornaments (*Pb/pb*). Orange converted to pinkish-purple by Pb. Asian Blau removes orange and Ivory removes yellow-orange revealing modified violet-blue iridophores and white leucophores. **(B)** Red Grass Delta expressing orange ornaments (*pb/pb*). Orange converted to pinkish-purple by Pb, photos courtesy of Taketoshi Sue. **(C)** Pink Grass Delta expressing Pb modified ornaments (*Pb/pb*). Orange converted to pinkish-purple by Pb, photo courtesy of Gyula Pasaréti.

**Figure 33.**
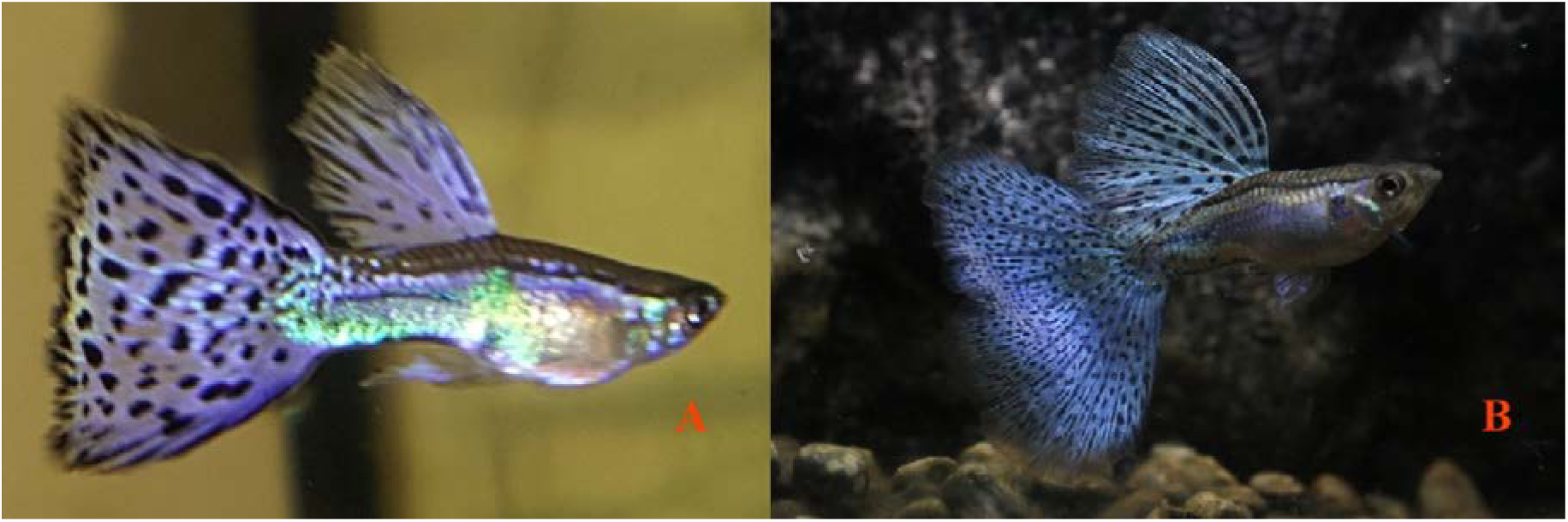
**(A)** Asian Blau (*Ab*) Purple Grass Delta expressing Pb modified ornaments (*Pb/pb*). Orange converted to pinkish-purple by Pb. Asian Blau removes orange revealing modified violet-blue iridophores. Photo courtesy of Leanne Shore. **(B)** Asian Blau (*Ab*) Blue Grass Delta (*pb/pb*). Orange converted to pinkish-purple by Pb. Asian Blau removes orange revealing modified blue iridophores. Photo courtesy of Kevin Yao.

**Figure 34.**
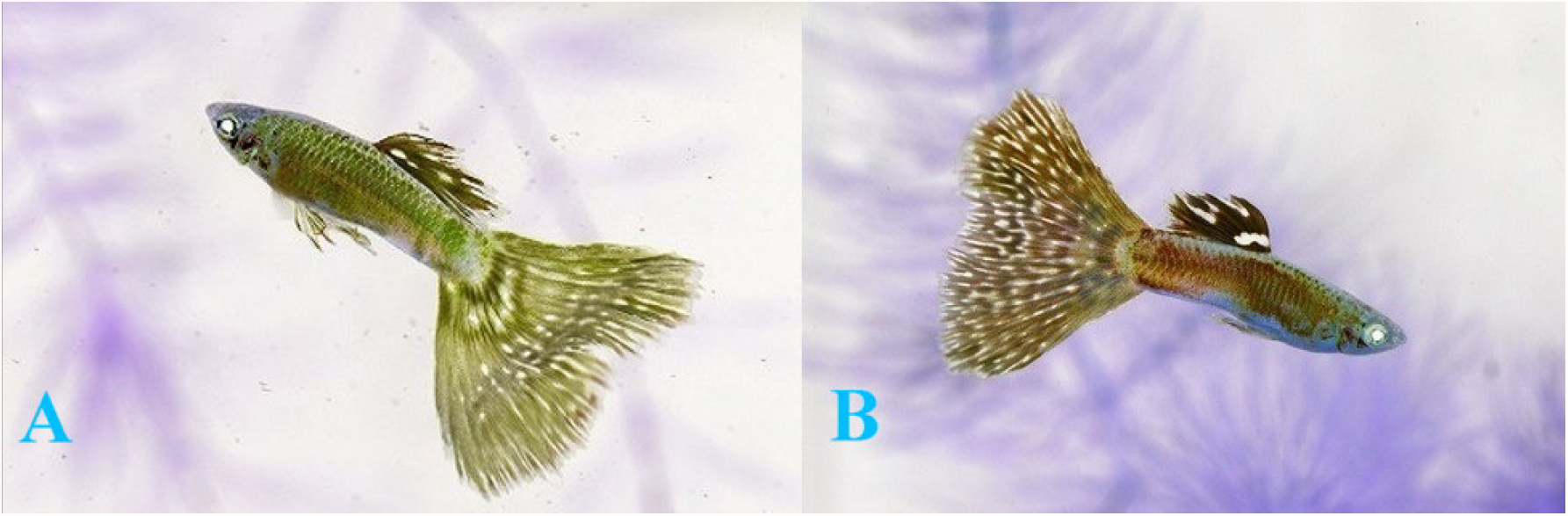
**(A)** Asian Blau (*Ab*) Ivory (i); i.e. Lavender Coral Red (*Co*) Grass Delta expressing Pb modified ornaments (*Pb/pb*). Orange converted to pinkish-purple by Pb. Asian Blau removes orange and Ivory removes yellow-orange revealing modified violet-blue iridophores and white leucophores. **(B)** Asian Blau (*Ab*) Coral Red (**Co**) Blue Grass Delta (*pb/pb*). Orange is converted to pinkish-purple by Pb. Asian Blau removes orange revealing modified blue iridophores and minimal Metal Gold (**Mg**). Photos courtesy of Taketoshi Sue.

**Fig 35.**
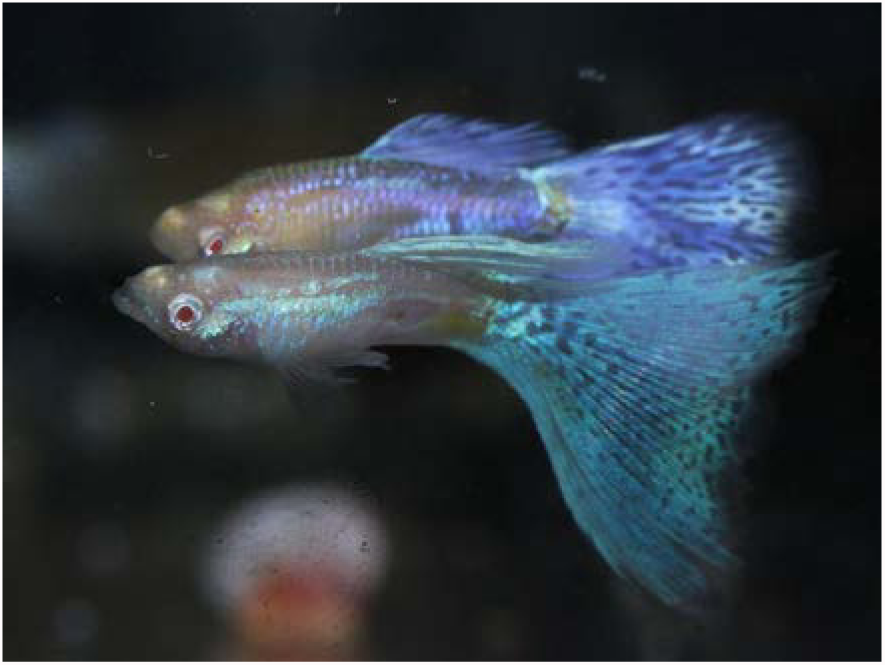
**Top:** Asian Blau (*Ab*) Albino (*aa*) Ivory (*ii*); i.e. Albino Lavender Grass Delta expressing Pb modified ornaments (*Pb/Pb or Pb/pb*). Albino removes melanophores. Orange converted to pinkish-purple by Pb. Asian Blau removes orange and Ivory removes yellow-orange revealing modified violet-blue iridophores. **Bottom:** Asian Blau (*Ab*) Albino (*aa);* i.e. Albino Blue Grass Delta. Albino (*aa*) removes melanophores. Asian Blau (*Ab*) removes orange revealing modified blue iridophores and Metal Gold (*Mg*). Photos courtesy of Taketoshi Sue.

**Fige 36.**
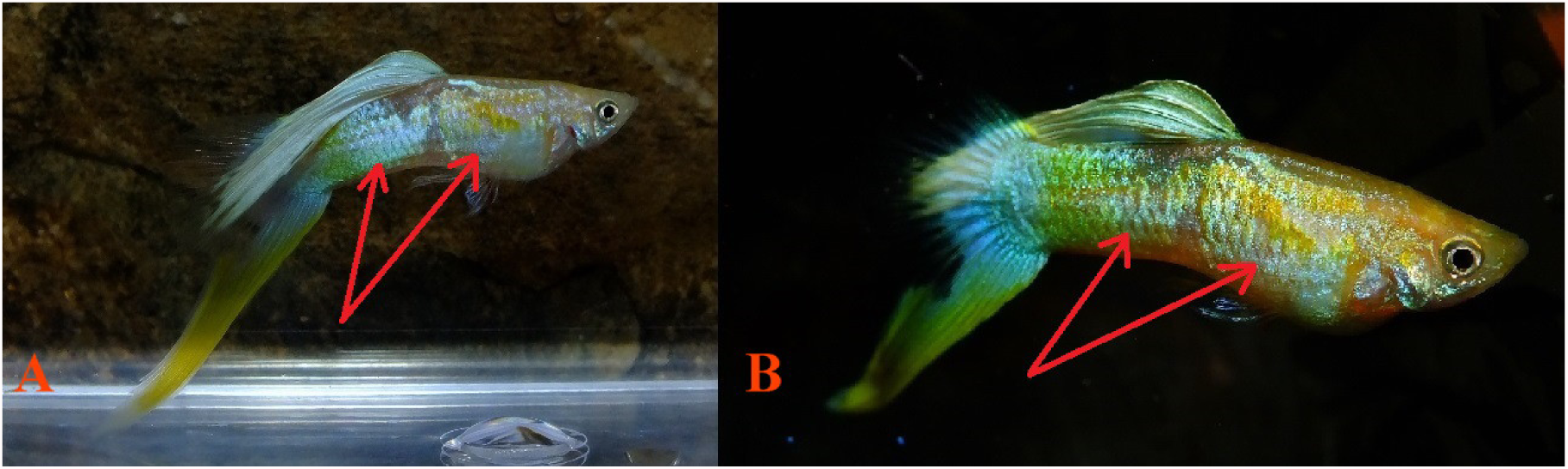
**(A)** Asian Blau (*Ab*) Blond (*bb*) Vienna Lowersword expressing Pb modified ornaments (*Pb/Pb*). **(B)** Asian Blau (*Ab*) Blond (*bb*) Vienna Lowersword (*Pb/pb*). Blond reduces melanophores. Orange converted to pinkish-purple by Pb, by xanthophore removal (arrows). Asian Blau removes orange revealing modified light violet-blue iridophores.

## Discussion and Conclusions

The genes kita, kitla, csf1ra, and csf1rb have been identified in Guppies (Kottler 2013; 2015).

Kita is a required precursor gene in the development of early forming motile melanophores (kita-dependent) in Guppies. Kita is not required for development of genetically distinct late forming, possibly non-motile, melanophores (kita-independent). Together, they produce the reticulated scale pattern found in both sexes; wild and domestic.

Csf1ra is a required precursor gene (colony-stimulating factor 1 receptor) in the development and formation of xantho-erythrophores. In Guppies csf1ra is not required in development of late forming, possibly non-motile, melanophores (kita-independent). Together, they contribute to non-defective ornaments of wild-type males.

Interactions between late forming melanophores in conjunction with xantho-erythrophores affect Domestic Guppy strain ornaments in both sexes. Thus, functional kita and csf1ra are found in non-defective ornaments of Domestic strains in both sexes.

Purple body removes certain classes of yellow-orange-red color pigment over silver iridophores or white leucophores. Dark red pteridine color pigment does not seem to be modified by Purple Body in fins lacking an underlying silver iridophore or white leucophore pattern. Modification by Pb seems limited predominantly to wild-type orange color pigmented xantho-erythrophores; i.e. those which also contain yellow-orange carotenoids in addition to red pteridines, over an iridophore pattern. Heterozygous Purple Body (*Pb/pb*) alters orange spots in select regions of the body and in finnage to “pinkish-purple”.

Heterozygous Pb does not appear to greatly reduce visible structural yellow color pigment cells over white leucophore or reflective clustered yellow cells in body and finnage. A slight increase in visibility of violet and blue iridophores is often found. Homozygous Pb expression results in a further removal of xantho-erythrophores, in conjunction with both increased populations and/or greater visibility of modified melanophores and naturally occurring violet and blue iridophores. Pb/Pb plus an unidentified additional genetic component is required for production of the all-purple phenotype.

When Pb is combined with any of the autosomal or sex-linked mutants well known to breeders, Albino (*aa*), Blond (*bb*), Golden (*gg*), and Asian Blau (*Ab*), further combined effects occur. The modifications of each of these genes on “wild-type” grey and resulting co-expressions with the Purple Body (*Pb*) gene are briefly discussed as follows:

## Autosomal Genes With References

### Albino

(*aa*). Recessive. Also known as Real Red Eye Albino (*RREA*) or (*Type* B). There is an inability to produce black melanophores in the body and finnage. It eliminates all classes of melanophores; dendritic, corolla and punctate. Therefore melanin is absent as well. Albino is epistatic to both the Blond and Golden genes; thus mutant alleles of each may be found in the albino genotype (*e.g., aa Bb, aa bb, aa Gg, aa gg, etc*.). In Albino, the result when combined with Pb is similar to that with Blond, except that the melanin is completely absent and the colors appear even paler (except when a different pigment cell is over expressed as in full red). Red can appear deepened and “darker” when Pb is present.

### Blond

(*bb*). Recessive. It has been identified as a defective gene mutation in adenylate (adenylyl) cyclase 5 (adcy5; ac5), and mapped to Linkage Group (LG) 2 (Kottler et al. 2015). It is also known to pedigree breeders as IFGA Gold, Asian Gold, European Blond. This mutation produces a near normal number of black melanophores of all types; Dendritic, Corolla and Punctate. However, the size of each is greatly reduced and the structure modified as compared to “wild-type” grey. According to published results Blond is not linked to Golden. This gene should be viewed not as a suppressor of melanophores, but rather one that alters melanophore size and shape.

When Pb is combined with homozygous blond (*bb*), a violet-blue sheen co-expresses with often paler and less intense xantho-erythrophores, resulting from the reduction in melanophore size. The reflective qualities of xantho-erythrophores in Blond are determined by the composition of underlying structural colors (iridophores are more reflective and leucophores are less reflective) and angles at which crystalline platelets reside.

### Golden

(*gg*). Recessive. The Golden gene is a defective ortholog of kita (Kottler et al. 2013). Kita has been mapped on Guppy autosomal LG 4 (Tripathi et al. 2008). It is also known to pedigree breeders as European Gold, IFGA Bronze or Asian Tiger. This gene produces a reduced amount of black melanophores (approximately *50*%) of all types; dendritic, corolla and punctate. However, the size of dendritic and corolla melanophores is greatly increased and the collection of corolla melanophores is concentrated into “clumps” or “islands” of melanophores along the scale edges. Males and females lack skin melanophores at birth, but they develop with maturity. Scale edging will become lighter with a higher inbreeding co-efficient; i.e. long-term Golden x Golden breeding’s. This suggests that there are non-allelic modifier genes affecting the Golden phenotype.

According to published results Golden and Blond are in different linkage groups and assumedly different chromosomes and thus *can not* be alleles of each other or linked to each other. This gene should be viewed as a suppressor of melanophore population numbers.

In Pb plus Golden (*gg*), the effect is similar to Blond, except that melanophores are present at higher frequencies and with a modified distribution. As a result, the colors while paler than in grey are not as pale as in Blond.

### Asian Blau

(*Ab*). Incompletely Dominant. Also known to pedigree breeders as (*r2*) Europe and (*Rr*) Asia. **[Note:** The use of lower case “r” violates the accepted genetic use of symbols since this is not a recessive gene, this usage came about prior to identification of Ab as a second erythrophore defect.] In heterozygous condition red color pigment is removed, while collected yellow color pigment and clustered Metal Gold (*Mg*) is little affected. This produces an iridophore based phenotype. Snakeskin patterns degrade in both heterozygous and homozygous expression, as a result of disruption of melanophore structure or melanin content. The Purple (Violet) sheen found above the lateral line of both males and females is removed.

In homozygous condition certain black melanophores are removed along with red and yellow color pigments. In homozygous condition finnage may be reduced in size, but the genes are still present in the genotype for normal finnage. An outcross of homozygous Ab will produce the expected finnage in F_1_ offspring. **[Note:** As there are distinct types of red color pigment (carotenoid and pteridine) present in both body and finnage, removal may not be complete, as in a red “Old Fashioned” shoulder stripe. A very faint “red shoulder stripe” is sometimes visible.]

The result when Asian Blau is combined with Pb can range from highly reflective violet-blue to non-reflective violet-blue in combination with additional genes that remove and/or reduce iridophores or alter angles of crystalline platelets.

### European Blau

(*r* or *r1*); also (*eb*). Recessive. Both csf1ra and csf1rb genes have been identified in Guppies, and are the result of an ancestral genome duplication event that produced four copies of each gene rather than two. (The guppy is an ancestral tetraploid.) In many fish species one or the other pair of some genes has been lost to reduce the total gene dosage back to a “diploid level” of two rather than four copies. In some other cases, the two genes diverge from each other and assume different functions. The European Blau gene is a defective ortholog of csf1ra. Expression levels of csf1rb were not upregulated to compensate for the deficiency in csf1ra (Kottler et al., 2013), which suggests that Csf1ra and Csf1rb have functionally diverged from each other in the guppy. Csf1ra has been mapped on Guppy autosomal LG 10 (Tripathi et al. 2008).

European Blau is also known as Dunkel in Asia. Csf1ra activity is required for the dispersal or differentiation of male-specific xanthophores (Kottler et al, 2013). In homozygous condition it is epistatic to wild type genes for red and yellow; major red and yellow color pigments are removed from the body. Certain red color pigments may be present in finnage, and to a lesser degree in the body. Reflective qualities are reduced. Ecotopic melanophores may be removed, while basal level melanophores such as are found in Half Black (NiII) are only slightly reduced. “The salient feature of the csf1ra mutant males was the absence of all orange traits, with concomitant severe changes in black ornaments” (Kottler et al, 2013). Snakeskin patterns degrade in homozygous expression. The purple (violet) sheen found above the lateral line of both males and females is removed. There is minimal finnage reduction.

## Additional Autosomal Genes Referred To

### Pink

(p, Luckman 1990; Förster 1993; *pi* Kempkes 2007) Recessive. Removal of orange erythrophores in body resulting in a “yellow-orange” cast in finnage. Homozygous reduction of NiII melanophores and increase in MBAG. Removes blue iridophores. Reduces size of finnage. Pb modification: Orange spotting is converted to pinkish-purple. Collected yellow pigment cells are removed, but not Clustered Mg. Body color may be modified to violet-blue.

### Ivory

(*I*, Tsutsui, Y 1997) Autosomal Dominant. Heterozygous suppression of erythrophores (red color). Homozygous suppression of xantho-erythrophores (yellow-red color), with reduction in fin size. Possible differences in melanophore modifications in heterozygous vs homozygous states. Resulting in a “white” appearance. II (homozygous), Ii (heterozygous) and ii (non-Ivory). Pb modification: Previous orange spotting is removed by Ivory. Underlying iridophores and leucophores converted to light pinkish-purple.

### Magenta

(*M, undescribed*) Autosomal Dominant. Proliferation of red color pigment when present and an increase of violet-blue iridophore structural color. Converts yellow color pigment cells (xanthophores) to red erythophores), though Metal Gold (*Mg*) may remain. Concentrates black melanophores. There is a reduction in fin size. Pb modification: Orange spotting is converted to pinkish-purple. Collected yellow pigment cells are removed, but not Clustered Mg.. Converts orange to red and deepens violet blue coloration.

### Zebrinus

(*Ze*, Winge 1927) Autosomal Dominant. Color Character; Barred pattern of vertical stripes on the peduncle, viz. 2-5, generally 3 dark pigment stripes. Effect resembles that of Tigrinus gene, but is as a rule more pronounced. ZeZe (homozygous), Zeze (heterozygous) and zeze (non-Zebrinus). Pb modification: No direct effect on barring pattern. Overlaying orange spotting is converted to pinkish-purple. Collected yellow pigment cells are removed, but not Clustered Mg.

## Major Sex-linked Traits Referenced

### Coral Red

(*Co, undescribed*) Y-linked. Red color pigment shoulder pattern. Linked in complex with Ds. Probably a Full Body modifier. It originated out of Vienna Emerald Green Ds. Pb modification: Proliferation of violet structural color. Orange spotting is converted to pinkish-purple. Collected yellow pigment cells are removed, but not clustered Mg.

### Grass

(*Gra*, Tsutsui, Y. 1997; Iwaski, N. 1989) X and/or Y-linked dominant. The Grass phenotype is a highly variable random “fine dot” circular melanophore pattern in finnage. Primarily limited to caudal ornamentation, with limited dorsal influence. Often associated with “Nike Melanophore Stripe” body pattern. Variegation shape is dependent upon in-breeding co-efficient. Color pigments can be added. “Glass Grass” genotype is similar to Multi with a translucent background and color pigments. “Grass Grass” genotype is often linked in complex with sex-linked xantho-erythrophore color pigment spots. Pb Modification: Some orange spotting is converted to pinkish-purple. Collected yellow pigment cells are removed, but not Clustered Mg. Xanthophore removal may reveal white leucophores if present.

### Moscow Blau Additional Gene

(*MBAG, undescribed*) X-linked dominant. Half body pattern expressing motile black mediating moderate & translucent melanin development over entire body area posterior to dorsal fin, and in caudal peduncle. Other posterior peduncle color patterns may be nearly or wholly obscured. Early Russian MBAG strains, in addition to NiII, may have been identified as “Tuxedo; i.e. HalfBlack” (Pg. 58, Iwasaki 1989). Pb modification: Orange spotting is converted to pinkish-purple. Collected yellow pigment cells are removed, but not Clustered Mg. Peduncle may take on violet-blue reflective coloration.

### Mosaic

(Mo, Khoo and Phang 1999b) X-linked dominant. The Mosaic phenotype is a highly variable random “large spot” crescent shaped melanophore pattern in finnage. Primarily limited to caudal ornamentation, with limited dorsal influence. Variegation shape is dependent upon in-breeding co-efficient. Normally associated with erythrophore color pigment (carotenoid and/or pteridine). Pb modification: Some orange spotting in the body is converted to pinkish-purple. Collected yellow pigment cells are removed, but not Clustered Mg. Xanthophore removal may reveal white leucophores if present. Dark Red Caudal pigment is generally not modified.

### Multi

(–, *undescribed*) X-linked dominant. The Multi phenotype is a highly variable random “fine dot” circular melanophore pattern in finnage. Primarily limited to caudal ornamentation, with limited dorsal influence. Variegation shape is dependent upon in-breeding co-efficient. Color pigments can be added. Not linked with erythrophore color pigment (carotenoid and/or pteridine) spots. This must be added through outcrossing. Little or no effect on existing body color or pattern. Pb modification: Some orange spotting in the body is converted to pinkish-purple. Collected yellow pigment cells are removed, but not Clustered Mg. Xanthophore removal may reveal white leucophores if present. Dark Red Caudal pigment is generally not modified.

### Moscow

(*Mw*, Kempkes 2007) Y-linked. Blue iridophore shoulder pattern. Likely a Full Body modifier. Color variation with addition or removal of xantho-erythrophores. Pb modification: Orange spotting is converted to pinkish-purple. Collected yellow pigment cells are removed, but not Clustered Mg. Body color is modified to violet-blue.

### Variegation

(*Var*, Khoo and Phang 1999). (See Grass, Mosaic, Multi) X and / or Y-linked dominant gene. Inheritance of variegated tail patterns appears to be determined by a single locus on the X and Y chromosomes.

The gene study of Variegation focused on variable random “large spot” shaped melanophore pattern in the caudal, though specimens exhibited similar dorsal pattern. Variegation shape is dependent upon in-breeding co-efficient. Color pigments (xantho-erythrophores) were not linked. Pb modification: Some orange spotting in the body is converted to pinkish-purple. Collected yellow pigment cells are removed, but not Clustered Mg. Xanthophore removal may reveal white leucophores if present. Dark Red Caudal pigment is generally not modified.

### Nigrocaudautus

(*NiI*, Nybelin 1947 and NiII, Dzwillo 1959) X and/or Y-linked dominant gene. Full body modifier, epistatic to many other genes in outcrosses. Pb modification: Orange spotting is converted to pinkish-purple. Collected yellow pigment cells are removed, but not Clustered Mg.

### Schimmelpennig Platinum

(*Sc*); **Buxeus** (Kempkes 2007). Y-linked dominant gene. Silver-Blue iridophore shoulder pattern with Metal Gold (**Mg**) overlay. Probably a Full Body modifier. Linked in complex with Ds. Originated out of Vienna Emerald Green Ds. Pb modification: Orange spotting is converted to pinkish-purple. Collected yellow pigment cells are removed, but not Clustered Mg.

## SUMMARY

The newly described gene Purple Body and prior described genes Albino, Blond, Golden, Asian Blau, and European Blau each limits or otherwise reduces the normal expression of chromatophores found in “wild-type” Grey. When they are combined together and with other frequently used color genes, new phenotypes are produced which are useful to Pedigree Guppy Breeders and Commercial Farmers alike. Their basic effects are of interest to geneticists, biochemists and molecular biologists.

### Photo Imaging

Photos by author(s) were taken with a Fujifilm FinePix HS25EXR; settings Macro, AF: center, Auto Focus: continuous, varying Exposure Compensation, Image Size 16:9, Image Quality: Fine, ISO: 200, Film Simulation: Astia/Soft, White Balance: 0, Tone: STD, Dynamic Range: 200, Sharpness: STD, Noise Reduction: High, Intelligent Sharpness: On. Lens: Fujinon 30x Optical Zoom. Flash: External mounted EF-42 Slave Flash; settings at EV: 0.0, 35mm, PR1/1, Flash: -2/3. Photos cropped or brightness adjusted when needed with Microsoft Office 2010 Picture Manager and Adobe Photoshop CS5. All photos by author(s), unless otherwise noted.

### Ethics Statement

No specimens were euthanized or harmed in this study.

### Notes

This publication is number three (3) of four (4) by Bias and Squire in the study of Purple Body (*Pb*) in *Poecilia reticulata:*

1. The Cellular Expression and Genetics of an Established Polymorphism in *Poecilia reticulata;* “Purple Body, (Pb)” is an Autosomal Dominant Gene,
2. The Cellular Expression and Genetics of Purple Body (*Pb*) in *Poecilia reticulata*, and its Interactions with Asian Blau (*Ab*) and Blond (*bb*) under Reflected and Transmitted Light,
3. The Cellular Expression and Genetics of Purple Body (*Pb*) in the Ocular Media of the Guppy *Poecilia reticulata*,
4. The Phenotypic Expression of Purple Body (*Pb*) in Domestic Guppy Strains of *Poecilia reticulata*.

## Acknowledgements

To my best friend and wife Deana Bias, for her support and persistence over the last several years in this four part study… To my co-author and dear friend Rick Squire for his patience as a mentor… To those Domestic Breeders who willingly provided additionally needed pedigree strains and study populations for completion of this paper…

